# PIEZO1 mediates mechanosensing and myofibroblastic activation of synovial fibroblasts in post-traumatic osteoarthritis

**DOI:** 10.64898/2026.10.13.758946

**Authors:** Easton C. Farrell, Aanya Mohan, Alexandra Kahn, Eleanor Knight, Hunter Stabile, Morgan Whitaker, Sung Yeon Kim, Lindsey Lammlin, Michael D. Newton, George Esteve, Ranveer R. Singh, Scarlet C. Howser, Selina Z. Lin, Huong X. Tran, Danny M. Nguyen, Kajetana Bevc, Sofia Gonzalez-Nolde, Lin Han, Robert L. Mauck, Carla R. Scanzello, Kurt D. Hankenson, Alexander J. Knights, Tristan Maerz

## Abstract

Fibrosis and inflammation of the synovium are common pathological features of joint diseases. In osteoarthritis (OA), extensive evidence supports that synovial pathology drives symptoms and disease progression, yet the mechanisms underpinning pathological synovial remodeling, including fibrosis, remain unclear. Here, we identify the mechanosensitive ion channel PIEZO1 as a critical regulator of myofibroblast-like activation of synovial fibroblasts (SFs) in post-traumatic osteoarthritis (PTOA). Using multi-omic assessments in human and murine synovium, a murine model of joint injury, histopathology, and functional *in vitro* phenotyping, we demonstrate that joint injury triggers expansion of myofibroblast-like SFs concurrent with fibrotic remodeling and mechanical stiffening of the synovium. Myofibroblastic commitment of synovial progenitors was associated with activation of the YAP1 regulon, and PIEZO1 agonism drove nuclear accumulation of YAP1 in SFs. PIEZO1 was enriched in synovia of injured joints, and in cultured SFs pharmacologic PIEZO1 activation promoted αSMA enrichment and increased matrix contraction, which were both ablated by YAP1 inhibition. PIEZO1 silencing abrogated TGF-β-induced contractility and markedly altered the transcriptomic response to mechanical loading *in vitro*, including ECM production and neuroinflammatory programs. *In vivo*, acute intra-articular pharmacologic PIEZO1 activation induced a pro-fibrotic, pro-inflammatory, and anti-adipogenic transcriptomic program in synovium. Collectively, these findings establish PIEZO1 as a key mediator of the pathogenic activation of SFs and identify aberrant mechanosensing as a mechanism underlying synovial fibrosis in PTOA.

## Introduction

Fibrosis and inflammation of the synovium – the inner soft tissue lining of joints – are hallmarks of osteoarthritis (OA), a debilitating and prevalent disease marked by pain, stiffness, and progressive loss of joint function^1–3^. Although historically considered a disease of cartilage and bone, OA is now recognized to also be driven by synovial pathology. Fibrosis and inflammation converge and are clinically associated with pain in multiple diseases, including OA^3–7^. However, effective anti-fibrotic therapies are lacking, and no disease-modifying OA treatments exist beyond palliative analgesic interventions or joint replacement. Development of new effective OA treatments depends on defining the largely unclear mechanisms of pathological tissue remodeling, including progressive synovial fibrosis.

Single-cell studies have revealed the heterogeneous and dynamic cellular landscape of human and murine OA synovium^8–12^. These revealed disease-associated synovial fibroblasts (SFs) with diverse pathological functions spanning inflammation, extracellular matrix (ECM) remodeling, and neurotrophic activity. Disease-enriched subsets include cells resembling myofibroblasts, which express markers of fibrotic progression. While critical in wound repair, myofibroblast persistence drives fibrosis and tissue dysfunction in numerous contexts^13^. In end-stage human OA synovium, advanced fibrosis and enrichment of myofibroblast-like cells have been reported^14^, and *LRRC15*-expressing myofibroblast-like SFs were enriched in patients reporting high pain at arthroplasty^12^.

Joint injury-induced OA, termed post-traumatic osteoarthritis (PTOA), triggers myofibroblast-like SF emergence concurrent with an incompletely resolving inflammatory response, as demonstrated in murine injury models. Using surgical destabilization of the medial meniscus (DMM) and non-invasive anterior cruciate ligament rupture (ACLR), we and others have shown rapid synovial enrichment of myofibroblast-like cells expressing canonical markers such as *Acta2*, *Tagln, Comp, and Lrrc15,* which persist through OA development^8,11,15^.

Despite the pathological relevance of myofibroblast-like SFs in PTOA, their disease-associated spatiotemporal dynamics and the mechanisms driving their persistence are not fully understood. We previously used scRNA-seq trajectory modeling to predict that synovial *Pi16+ Dpp4+* stromal cells, transcriptionally similar to the “universal fibroblast”^16^, could serve as progenitors for pathological SF subtypes, including *Prg4-hi* lining SFs via an *Acta2*/αSMA+ myofibroblast-like intermediate^8^. A similar trajectory was identified in the DMM model^11^, confirmed by lineage tracing^15^. In homeostasis, *Pi16+ Dpp4+* fibroblasts function as adipogenic progenitors giving rise to adipocytes in niches including intra-articular adipose tissue^15^. Dysregulation of their fate commitment is a potential mechanism of fibrosis, and the upstream activators governing this transition require further attention.

Aberrant mechanosensing has gained interest as a potential activator of myofibroblastic differentiation, fibrogenic cells in chronic disease, and tissue fibrosis progression^17,18^. Overactivity of mechanoresponsive pro-fibrotic pathways may arise from abnormal loading, including micro-level alterations in ECM stiffness and macro-level changes in joint kinematics described in ACLR^19–21^. Across fibrosis contexts, the TGF-β-SMAD and canonical Wnt/β-Catenin (cWnt) signaling pathways have been linked to myofibroblastic activation^22–25^, and these pathways are mechanoregulated^26–29^. YAP1, a mechanosensitive transcriptional co-factor and main effector of the Hippo pathway, interacts with these pathways to promote myofibroblast activation in the heart, lungs, and kidneys^30–33^. YAP1 upregulation has been observed in SFs from both OA and rheumatoid arthritis (RA) synovium^34,35^, and YAP1-deficient mice exhibit reduced synovial hyperplasia in models of inflammatory arthritis and cartilage injury^36,37^. While mechanoresponsive, YAP1 cannot directly sense mechanical signals, and the upstream mechanoreceptors that activate it to perpetuate pro-fibrotic effects remain undescribed in synovial fibrosis.

This study dissected the mechanobiological mechanisms and spatiotemporal dynamics of myofibroblast-like SF emergence and persistence in PTOA synovium. Using an integrative multi-omics framework and functional studies at the cell and tissue level, we mapped myofibroblast-like SFs in the mechanically-stiffened synovium during disease. Trajectory modeling identified overactive YAP1 signaling governing the fate transition of synovial progenitors to myofibroblast-like SFs, and we described a functional PIEZO1–YAP1 axis driving fibrogenic SF loading responses to promote myofibroblastic activation and fibrotic remodeling.

## Results

### Synovial stiffening accompanies emergence of myofibroblast-like SFs in PTOA

We previously demonstrated post-injury phenotypic diversification of SFs, identifying emergent myofibroblast-like SFs with a pro-fibrotic, pro-inflammatory transcriptomic signature^8,38^. Extending this, we sought to identify the niches harboring myofibroblast-like SFs during PTOA progression. Using Xenium 5K Prime spatial transcriptomics, we generated a spatial atlas of SFs in the murine knee joint at true single-cell resolution (**Fig 1A-C**). Dimensionality reduction and semi-supervised clustering of anterior and posterior synovial compartments across n=12 histological sections (n=4 mice across uninjured (Sham), ACLR 7d, and ACLR 28d) identified 61,915 total cells (**Fig S1A**). Among 46,519 total SFs extracted and re-clustered independently (**Fig S1B**), we identified a synovial myofibroblast-like population expressing *Comp, Lum, Col8a1, Postn, Acta2, Tagln,* and *Sparc,* with pathway enrichment for ECM production, cytoskeletal contraction, TGFβ and Wnt signaling, neurotrophic activity, and inflammatory signaling (**Fig S1C, Supplemental Table 1 and 2**). Consistent with prior scRNA-seq studies of synovial cells,^8,11,12,38^ we further identified sublining fibroblasts, lining fibroblasts (*F13a1+ Clic5+*), progenitor fibroblasts (*Pi16+ Cd34+*), immune-interacting fibroblasts (*Vim+ Msn+*), and two subsets resembling osteochondral (OC) lineage cells (*Acan+ Sox9+*) (**Fig S1D, Supplemental Table 2**).

**Fig 1.**
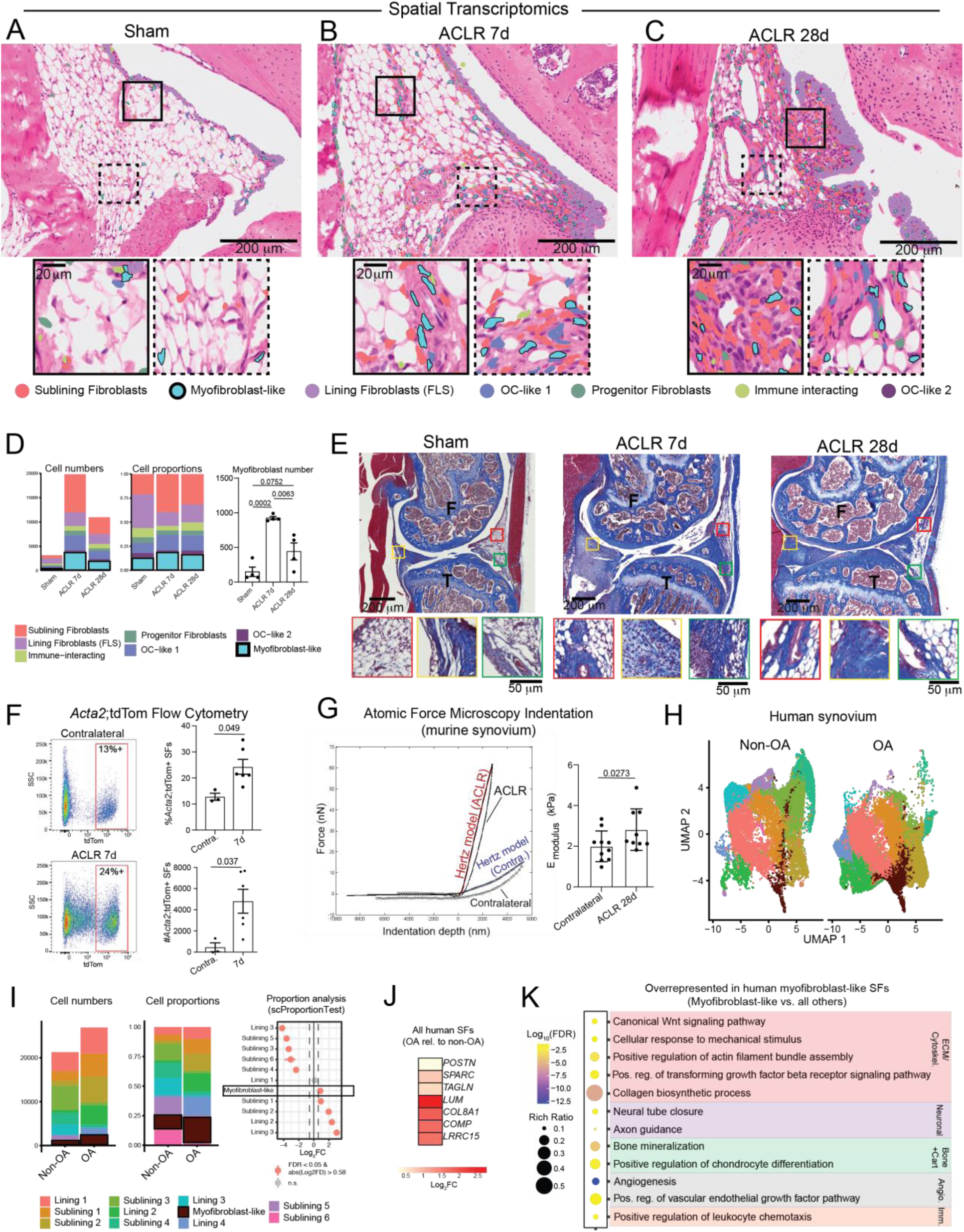
Synovial stiffening accompanies emergence of myofibroblast-like SFs in PTOA. **(A-C)** Xenium 5K spatial sequencing showing synovial fibroblast subset distribution in anterior synovium and IFP of Sham (A), ACLR 7d (B), and ACLR 28d (C) mice. Representative H&E-stained sagittal sections with overlays shown. Myofibroblast-like SFs outlined and highlighted in blue. High-magnification insets illustrate myofibroblast-like SFs in fibrotic regions. **(D)** Left/middle: number and proportion of total synovial fibroblast subsets identified in anterior and posterior synovium by spatial sequencing in Sham, ACLR 7d and ACLR 28d synovia. Right: Quantification of myofibroblast-like SFs across Sham, ACLR 7d and ACLR 28d synovium. n=4 (2 male, 2 female) per condition. One-way ANOVA with Tukey’s post hoc test. **(E)** Representative sagittal Masson’s Trichrome-stained sections of the medial joint compartment in Sham, 7d, and 28d mice. High-magnification insets illustrate fibrotic regions in the subsynovium and IFP. **(F)** Flow cytometry-based quantification of number and proportion of total tdTom+ cells from *Acta2*-Cre^ERT2^;tdTom (*Acta2*;tdTom) mice in ACLR 7d and contralateral synovia. n = 4 males and 4 females. Paired two-tailed t-test. Representative SSC vs tdTom fluorescence plots included (left). **(G)** Left: Representative force versus indentation depth plots from atomic force microscopy indentation of contralateral and ACLR 28d murine anterior synovium. Blue and red solid lines represent finite thickness-corrected Hertz model fitted to indentation data of contralateral and ACLR 28d, respectively. Right: Indentation moduli of contralateral and ACLR 28d murine anterior synovium. n = 10 mice (5 male, 5 female) per condition. Data represents averages of 15-20 indentations. Paired two-tailed t-test. **(H)** Uniform Manifold Approximation and Projection (UMAP) plots of human SFs from non-OA and OA subjects. **(I)** Left/middle: Total cell number and proportion of total human SF subsets in non-OA and OA groups. Right: scProportionTest quantification of relative enrichment of SF subsets between non-OA and OA subjects, with positive fold-change indicating enrichment in OA. **(J)** Differential expression of fibrosis-associated genes in OA vs non-OA SFs, with positive fold-change indicating enrichment in OA. **(K)** Selected Gene Ontology – Biological Processes (GO:BP) terms differentially enriched in human myofibroblast-like SFs compared to all other SFs, derived via statistical overrepresentation testing (P_adj_ < 0.05).

Myofibroblast-like SFs were rare Sham synovium, with only a few populating the synovial sublining and infrapatellar fat pad (IFP) **(Fig 1A, D**). In contrast, injured synovium at 7d and 28d post-ACLR showed marked enrichment of myofibroblast-like SFs in multiple synovial niches (**Fig 1B, C**), with 9.88- and 4.73-fold increased abundance relative to Sham, respectively (**Fig 1D**). In the anterior compartment, myofibroblast-like SFs localized primarily to the sublining, closely associated with dense connective tissue and perivascular regions. The IFP, a specialized intra-articular adipose depot that undergoes fibrotic remodeling after joint injury^39^, was also enriched for myofibroblast-like SFs. In posterior synovium of injured mice, abundant myofibroblasts occupied the sublining, associated with hypercellular fibrotic connective tissue (**Fig S1E**). Masson’s Trichrome staining of sagittal joint sections demonstrated marked increases in collagen-rich fibrotic tissue in the anatomic niches harboring myofibroblast-like SFs at 7d and 28d post-ACLR (**Fig 1E**). To orthogonally corroborate myofibroblast emergence after ACLR, we performed flow cytometry on synovium from *Acta2*-Cre^ERT2^;tdTom mice given tamoxifen at the time of joint injury. This revealed a dramatic increase in both the number and proportion of *Acta2*;tdTom+ synovial fibroblasts (gated as CD31-CD45-CD146-THY1+) in ACLR 7d compared to uninjured contralateral synovia, confirming expansion of myofibroblast-like SFs after injury (**Fig 1F, Fig S1F**).

Because the myofibroblastic phenotype is also associated with altered stiffness of the surrounding ECM, we performed atomic force microscopy (AFM)-based indentation testing on anterior synovia at the same anatomic location where myofibroblast emergence and fibrotic expansion were observed. This revealed a significant ∼50% increase in mean indentation modulus at 28d post-ACLR compared to contralateral synovium, a timepoint of PTOA with marked synovial fibrosis (**Fig 1G, Fig S1F**), confirming synovial ECM stiffening during PTOA progression.

To assess myofibroblast-like SFs in human OA synovium, we generated an integrated scRNA-seq atlas of n = 48,218 human non-OA and OA SFs from public datasets (**Fig 1H, Fig S2A-D, Supplemental Tables 3 and 4).** Myofibroblast-like SFs were proportionally enriched in human OA compared to non-OA synovium (**Fig 1I**). Collectively, SFs from OA synovium showed increased expression of pro-fibrotic, pro-inflammatory, and myofibroblast marker genes versus non-OA SFs (**Fig 1J**). Human myofibroblast-like SFs highly expressed markers such as *COL3A1*, *COMP*, *SPARC*, and *TNC* (**Fig S2E**) and overexpressed pathway terms for ECM production, actin filament assembly, neurotrophic activity, osteochondral processes, leukocyte chemotaxis, and angiogenesis (**Fig 1K, Supplemental Table 5**). Furthermore, human OA SFs exhibited enhanced TGF-β receptor signaling activity (**Fig 1K**), and both human OA and murine PTOA synovium showed marked upregulation of TGF-β signaling-related genes (**Fig S1G, H**), indicating enrichment of the canonical pathway activating myofibroblastic differentiation. Taken together, traumatic joint injury in mice induced synovial ECM stiffening, accompanied by emergence of myofibroblast-like SFs that occupy multiple synovial niches and transcriptionally resemble myofibroblast-like SFs identified in human OA synovium.

### PIEZO1-YAP signaling is activated in injury-induced myofibroblast-like SF emergence

Recent lineage-tracing and scRNA-seq studies showed that *Dpp4+/Pi16+* stromal progenitors – transcriptionally similar to the “universal fibroblast”^16^ – underpin emergence of both lining and myofibroblast-like SFs in PTOA synovium^8,15^. Using Monocle3-based trajectory analysis^41,42^ on our previously-published scRNA-seq data of murine SFs from Sham and ACLR synovia^8^, we identified a trajectory branch from *Dpp4+/Pi16+* progenitors towards either *Prg4*-hi lining fibroblasts or mature αSMA+ myofibroblasts (**Fig S3A**). We isolated the myofibroblast branch (**Fig 2A**) and confirmed increased expression of myofibroblast markers with increasing pseudotime (**Fig 2B**). To define potential mediators of myofibroblastic commitment of synovial progenitors, we identified genes positively correlated with pseudotime, then clustered them by expression covariance along the myofibroblast trajectory. This revealed two pseudotime-dependent gene modules enriched in the myofibroblast-like branch (**Fig S3B, Supplemental Table 6**). These gene modules were submitted to RcisTarget^43^ to identify transcription factor (TF) binding motifs overrepresented in their promoter regions. High-confidence motifs were associated with the TEAD, SMAD, and TCF TFs, which have established roles in canonical Wnt/β-catenin (cWnt) signaling, the TGF-β pathway, and myofibroblastic activation in other tissue and disease contexts^25,44–46^ (**Fig 2C, Supplemental Table 7**). These TFs were also upregulated in human OA SFs compared to non-OA (**Fig 2C**). Metascape TRUUST analysis, an independent method to predict molecular regulation by TFs, also identified enriched SMAD activity, plus RUNX2, SP1, and SP3 (**Fig S3C**), all also associated with myofibroblastic activation and tissue fibrosis^47,48^.

**Fig 2.**
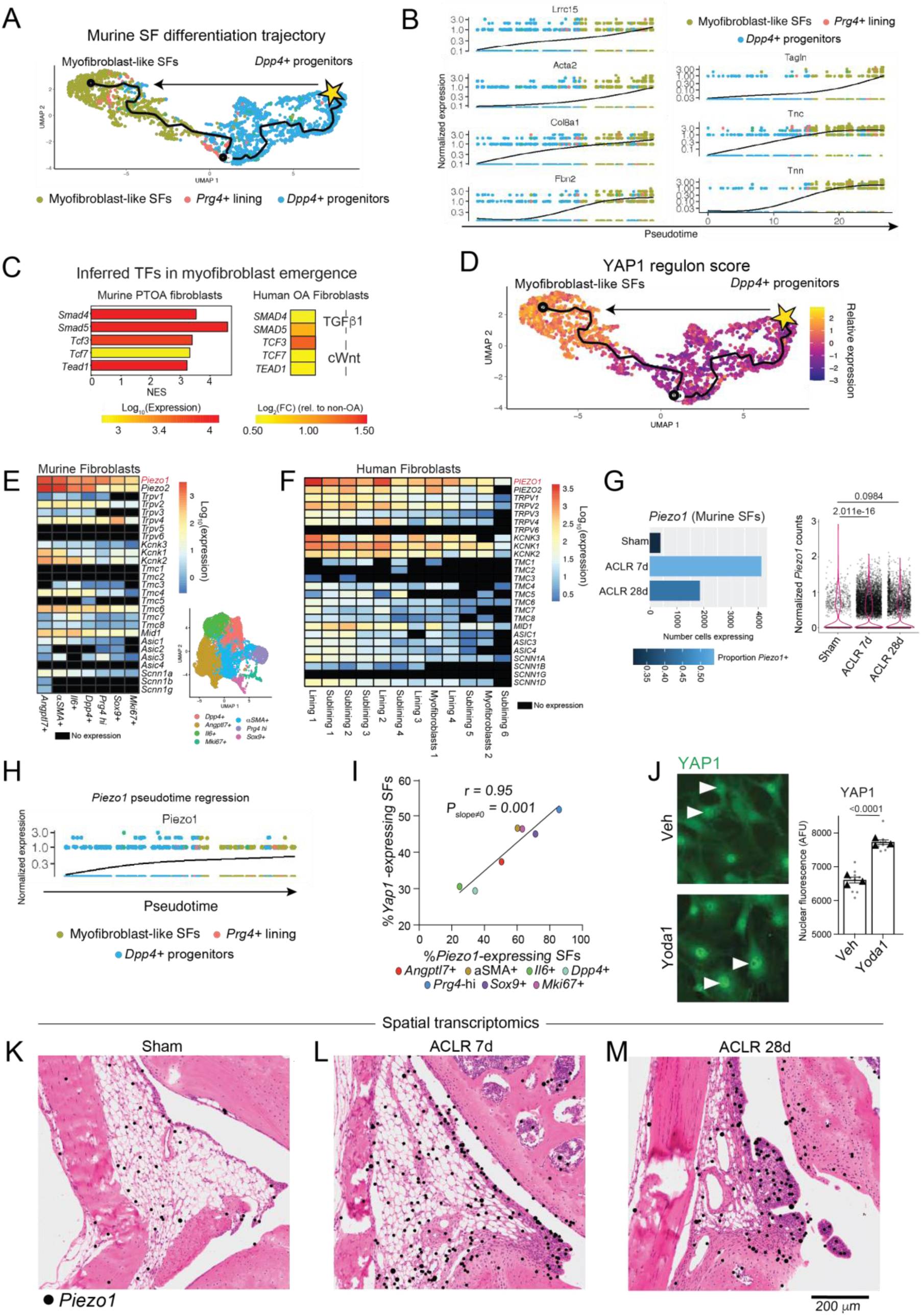
PIEZO1-YAP1 signaling is activated in injury-induced myofibroblast-like SF emergence. **(A)** UMAP plot of Monocle3-based cellular trajectory between murine *Dpp4*+ progenitor-like SFs and myofibroblast-like SFs. Trajectory was subset from the branching trajectory in Fig S3A. The star indicates the trajectory root node, and cells are colored by cluster annotation. **(B)** Pseudotime regression plots of myofibroblast-like SF marker genes along the differentiation trajectory from (A). **(C)** Left: Normalized enrichment scores (NES) of high-quality TFs (NES > 3, direct-annotation) predicted to regulate enriched pseudotime-dependent gene modules in the myofibroblast-like trajectory branch, derived via RCisTarget. Right: Differential expression of corresponding human genes, plotted as log_2_ fold-change in OA vs non-OA human SFs. **(D)** YAP1 regulon score representing aggregate relative expression of YAP1 target genes mapped onto the differentiation trajectory from (A). **(E)** Log-normalized gene expression of known mechanoreceptors and mechanosensitive factors across SF clusters in murine synovium (cluster annotation is shown in UMAP^8^), comprised of cells from Sham, ACLR 7d, and ACLR 28d. **(F)** Log-normalized gene expression of known mechanoreceptors and mechanosensitive factors across SF clusters from non-OA and OA subjects. Cluster annotation according to Fig S2B. **G)** Left: cell number and proportion of total SFs expressing *Piezo1* (normalized count >= 1) in murine synovia. Right: Violin plot of *Piezo1* expression in all murine SFs in Sham, ACLR 7d, and ACLR 28d. Wilcoxon rank sum test. **H)** Pseudotime regression plot of *Piezo1* expression along the differentiation trajectory from (A). **I)** Scatter plot of the percentage of *Piezo1*-expressing SFs vs the percentage of *Yap1*-expressing SFs. Seurat clusters according to Fig 2E. Pearson’s correlation coefficient and corresponding P value shown. Plotted line represents linear regression. **J)** Representative microscopy (left) and nuclear intensity quantification (right) of YAP1 immunofluorescent staining of murine SFs treated with vehicle or Yoda1 for 24h. n=3 technical replicate runs (each with 3 biological replicates pooled). Three fields of view per replicate. Two-tailed independent t-test. **(K-M)** Xenium 5K spatial transcriptomics showing spatial distribution of *Piezo1* transcripts (black dots) on representative H&E-stained sagittal sections of anterior synovium and IFP of Sham (K), ACLR 7d (L), and ACLR 28d (M) mice.

Mechanical cues such as loading and matrix stiffness are recognized to promote myofibroblastic activation^17,18,49^, prompting us to assess which mechanobiological axes may be active in myofibroblast-like SFs. Several of the implicated TFs identified in both murine and human SFs have known interactions with Yes-associated protein-1 (YAP1), a mechanosensitive transcriptional co-factor^50,51^. Relevant to canonical myofibroblast activation, YAP1 is a mechanoregulator of TGF-β-SMAD signaling and mediates crosstalk between the cWnt and Hippo pathways^52,53^. It has established roles in synovial hyperplasia, and YAP1-deficient mice demonstrate reduced synovitis in murine models of inflammatory arthritis and articular injury^36,37^. We therefore calculated YAP1 regulon activity from aggregate expression of known YAP1 target genes and confirmed it was highly enriched in the myofibroblast-like branch of the murine SF trajectory **(Fig 2D**). Together, these data implicate YAP1 as an effector of the transition from progenitor to myofibroblast-like SFs in PTOA synovia.

YAP1 is not itself a mechanoreceptor capable alone of transducing mechanical cues, though it does respond to upstream mechanoreceptors being activated. Given matrix stiffening in PTOA synovia and the predicted role of YAP1 in myofibroblast emergence, we next sought upstream mechanoreceptors that could sense synovial ECM modulus changes to activate YAP1 signaling and consequent myofibroblastic differentiation. Screening multiple families of mechanosensitive genes such as *Piezo*-, *Trpv*-, *Traak*- and *Trek-*family genes in SFs by scRNA-seq, we identified *Piezo1/PIEZO1* as the most highly expressed mechanoreceptor gene across all murine (**Fig 2E**) and human (**Fig 2F**) SF subsets. PIEZO1 is a mechanosensitive cation channel regulating Ca2+ signaling, and PIEZO1-mediated YAP1 activation was recently associated with myofibroblastic activation in pulmonary fibrosis^54^. *Piezo1* expression and the number of *Piezo1*-expressing SFs increased significantly in murine synovium following ACLR (**Fig 2G, Fig S3D**). *Piezo1* expression increased over pseudotime as murine myofibroblast-like SFs emerged (**Fig 2H**), and *PIEZO1* was more highly expressed in human OA SFs compared to non-OA SFs (**Fig S3E**). Across murine SF subsets, *Piezo1* and *Yap1* expression was highly correlated (r = 0.95, p-value = 0.001) (**Fig 2I**), and multiple established intracellular mediators of cation flux potentially downstream of PIEZO1 were upregulated in murine and human OA SFs (**Fig S3F**). Expectedly, *Piezo1* was also broadly expressed across other synovial cells such as macrophages and endothelial cells in both murine and human tissue (**Fig S3G**).

To establish a functional link between PIEZO1 and YAP1 in SFs, we treated primary murine SFs with Yoda1, a selective PIEZO1 agonist^55^, and observed significantly increased YAP1 nuclear protein content (**Fig 2J**). To assess whether SF YAP1 is responsive to increasing substrate stiffness, relevant to the synovial stiffening we observed in injured synovia (**Fig 1G**)^11^, we cultured primary murine SFs on soft (1.5 kPa) versus stiff (tissue-culture plastic, ∼1 GPa) substrates and observed a large stiffness-dependent increase in YAP1 nuclear protein content (**Fig S3H**).

We next used our spatial transcriptomic atlas to define the organization of *Piezo1*-expressing synovial cells. In uninjured joints, sparse *Piezo1*-expressing synovial cells were identified in the lining and sublining (**Fig 2K**). Joint injury increased *Piezo1*-expressing synovial cells, with ACLR 7d joints exhibiting abundant *Piezo1* expression primarily in lining, sublining, and IFP-resident cells (7.57- and 31.26-fold increase in transcript count compared to Sham in anterior and posterior synovia, respectively) (**Fig 2L, Fig S3I**). This pattern persisted to 28d post-ACLR, a timepoint of established PTOA marked by extensive synovial fibrosis (4.34- and 10.32-fold increase in transcript count compared to Sham in anterior and posterior synovia, respectively) (**Fig 2M, Fig S3I and J**). *Piezo1*+ myofibroblast-like SFs also increased significantly after injury (**Fig S3I**). As in our scRNA-seq analysis, spatial transcriptomics also identified broad *Piezo1* expression across multiple cell types and all SF subsets (**Fig S3K, L**). Collectively, these data implicate injury-induced PIEZO1-YAP signaling in mechanosensing of increased ECM stiffness and myofibroblast emergence in PTOA synovium.

### PIEZO1 mediates myofibroblastic activation of SFs in a YAP1-dependent manner

Given that PIEZO1 agonism induced YAP1 nuclear translocation in SFs (**Fig 2J**) and that trajectory analysis predicted YAP1 as a mediator of myofibroblastic fate commitment, we next tested the PIEZO1-YAP1 axis as a driver of myofibroblastic activation in SFs. Primary SFs from healthy murine synovium were treated with vehicle (DMSO), Yoda1 (10 µM), or the selective YAP1 inhibitor Verteporfin (0.5 µM), which blocks YAP1-TEAD binding and YAP1 nuclear translocation^56^ (**Fig S4A**). Immunofluorescent analysis demonstrated that PIEZO1 agonism significantly increased αSMA and F-actin protein, and concomitant YAP1 inhibition abrogated this αSMA increase (**Fig 3A**). αSMA localized to F-actin fibers, consistent with characteristic myofibroblast stress fiber enrichment (**Fig S4B**). Assessing SF migratory behavior, PIEZO1 agonism increased the capacity of SFs to fill a scratch wound, while YAP1 inhibition ablated this effect (**Fig 3B, Fig S4E**). Myofibroblasts also characteristically exert contractile force on their ECM. To determine whether PIEZO1-YAP1 activity affects this function, we employed a 3D collagen gel contraction assay and observed that PIEZO1 agonism significantly increased SF gel contraction, even in the absence of exogenous TGF-β, while YAP1 inhibition completely ablated this (**Fig 3C, Fig S4G**).

**Fig 3.**
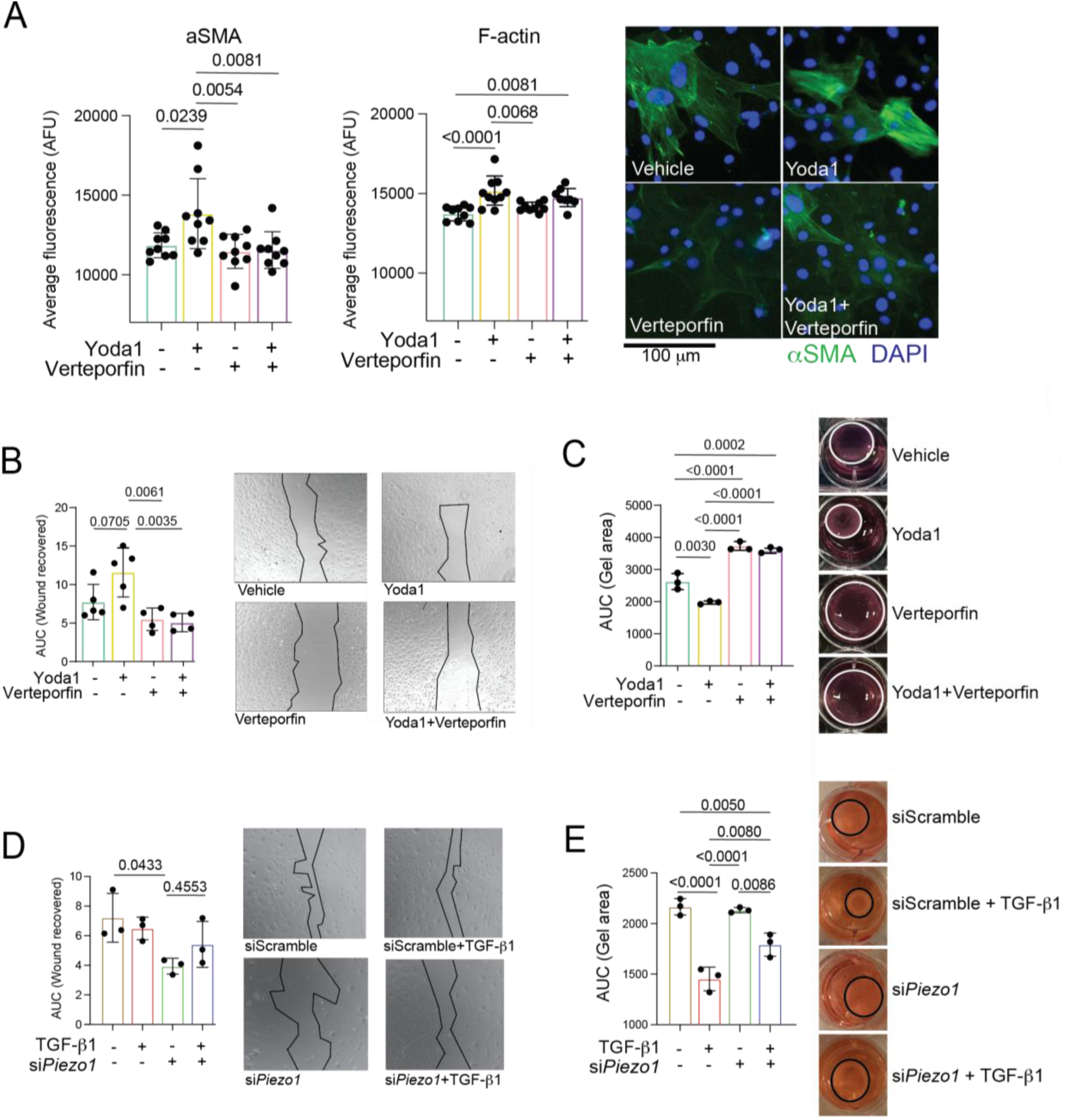
PIEZO1 mediates myofibroblastic activation of SFs in a YAP1-dependent manner. **(A)** Whole-cell fluorescent intensity quantification of αSMA (left) and F-actin (middle) immunofluorescent staining of murine SFs treated with vehicle, Yoda1, Verteporfin, or Yoda1 + Verteporfin for 24h. n=3 technical replicate runs (each with n=6 biological replicates pooled), with three fields of view quantified per replicate. One-way ANOVA with Tukey post-hoc testing. Welch’s correction applied to αSMA data. Representative fluorescent images of αSMA are shown on the right, with nuclei labeled by DAPI. **(B)** Quantification (left) and representative brightfield microscopy (right) of scratch wound assays in SFs pre-treated with vehicle, Yoda1, Verteporfin, or Yoda1 + Verteporfin for 24h prior to scratch introduction. n=4-5 technical replicate runs (n=6 biological replicates pooled). One-way ANOVA with Tukey post-hoc testing. **(C)** Quantification (left) and representative images at 24h (right) of gel contraction assays in collagen-embedded SFs pre-treated with vehicle, Yoda1, Verteporfin, or Yoda1 + Verteporfin for 24h prior to gel release and imaging for 24h. n=3 technical replicates n=6 biological replicates pooled). Two-way ANOVA with Tukey post-hoc testing. **D)** Quantification (left) and representative brightfield microscopy (right) of scratch wound assays in SFs pre-transfected with siPiezo1 or siScramble followed by treatment with vehicle or TGF-β1 for 24h. n=3 technical replicates (n=6 biological replicates pooled). One-way ANOVA with Tukey post-hoc testing. **E)** Quantification (left) and representative images at 24h (right) of gel contraction assays in collagen-embedded SFs pre-transfected with siPiezo1 or siScramble followed by treatment with vehicle or TGF-β1 for 24h prior to gel release and imaging for 24h. n=3 technical replicates (n=6 biological replicates pooled). One-way ANOVA with Tukey post-hoc testing.

TGF-β signaling is the canonical activator of myofibroblastic differentiation^57,58^, and we observed enhanced TGF-β pathway activity in SFs from murine and human OA synovium. (**Fig S1H**). To assess the requirement of PIEZO1 in TGF-β-induced myofibroblastic activation, we treated SFs with *Piezo1*-targeting siRNA (si*Piezo1*) to ablate Piezo1 (**Fig S4C**) or non-targeting siScramble, followed by TGF-β1 (10 ng/mL). TGF-β1 expectedly increased αSMA and F-actin protein (**Fig S4D**) and induced greater collagen gel contraction (**Fig 3E**) but did not alter wound closure rate, perhaps due to culturing on high-stiffness TCP which could mechanically activate myofibroblast-like SFs (**Fig 3D**). *Piezo1* silencing abrogated ∼50% of TGF-β1-induced gel contraction but did not alter contraction under basal conditions (**Fig 3E, Fig S4H**). si*Piezo1*-treated SFs also showed decreased wound healing capacity versus siScramble (**Fig 3C, Fig S4F**), but si*Piezo1* did not ablate αSMA or F-actin protein under basal or TGF-β1-treated conditions in 2D culture (**Fig S4D**).

Relevant to the established role of cWnt signaling in myofibroblastic activation^25,59^ and our prior studies demonstrating cWnt signaling overactivation in ACLR synovia and SFs^8^, PIEZO1 agonism with Yoda1 increased nuclear β-catenin content in murine SFs (**Fig S4I**), albeit not significantly. Yoda1 also strongly activated SuperTopFlash reporter signal in HEK293 cells, indicative of increased β-catenin-associated TCF/LEF activity, which concomitant YAP1 inhibition abrogated by ∼80% (**Fig S4J)**. Treatment of SFs with the β-catenin inhibitor PKF118-310 markedly decreased *Piezo1* expression (**Fig S4K**), suggesting a bidirectional interaction between PIEZO1 signaling and cWnt activity in SFs may exist. Taken together, PIEZO1 agonism promotes myofibroblastic activation via YAP1, partially mediates the functional effects of TGF-β signaling, and enhances cWnt signaling in SFs.

### PIEZO1 regulates the mechanical loading response of SFs

In PTOA, SFs may experience acute loading during the injury event as well as chronic mechanical activation from increased matrix stiffness and altered joint kinematics during PTOA progression. To assess PIEZO1 involvement in the SF loading response, we subjected SFs to cyclic tension after siRNA-mediated *Piezo1* silencing. 24h after a 4h cyclical loading bout (10% strain, 0.5 Hz), we confirmed comparable cell viability between conditions (**Fig S5A**) and then harvested RNA for bulk RNA-seq (**Fig 4A)**. After quality control (**Supplemental Table 8**), differentially expressed genes (DEGs) and pathways were derived. Vehicle (no transfection) and siScramble conditions exhibited no significant transcriptomic differences (**Fig S5B-C**); thus, siScramble served as the baseline control. *Piezo1* silencing dramatically altered the SF transcriptome under both unloaded and loaded conditions (**Fig 4B, Supplemental Table 9**). In unloaded SFs, si*Piezo1* yielded 579 DEGs (p_adj_<0.05, |Log_2_FC|>0.585, FPKM>1) versus siScramble (**Fig S5C**), mapping to downregulated gene ontology terms for ECM maintenance, including bone and cartilage-related terms, cWnt signaling, inflammatory and neurotrophic activity, fat cell proliferation, and angiogenesis (**Fig 4C**).

**Fig 4.**
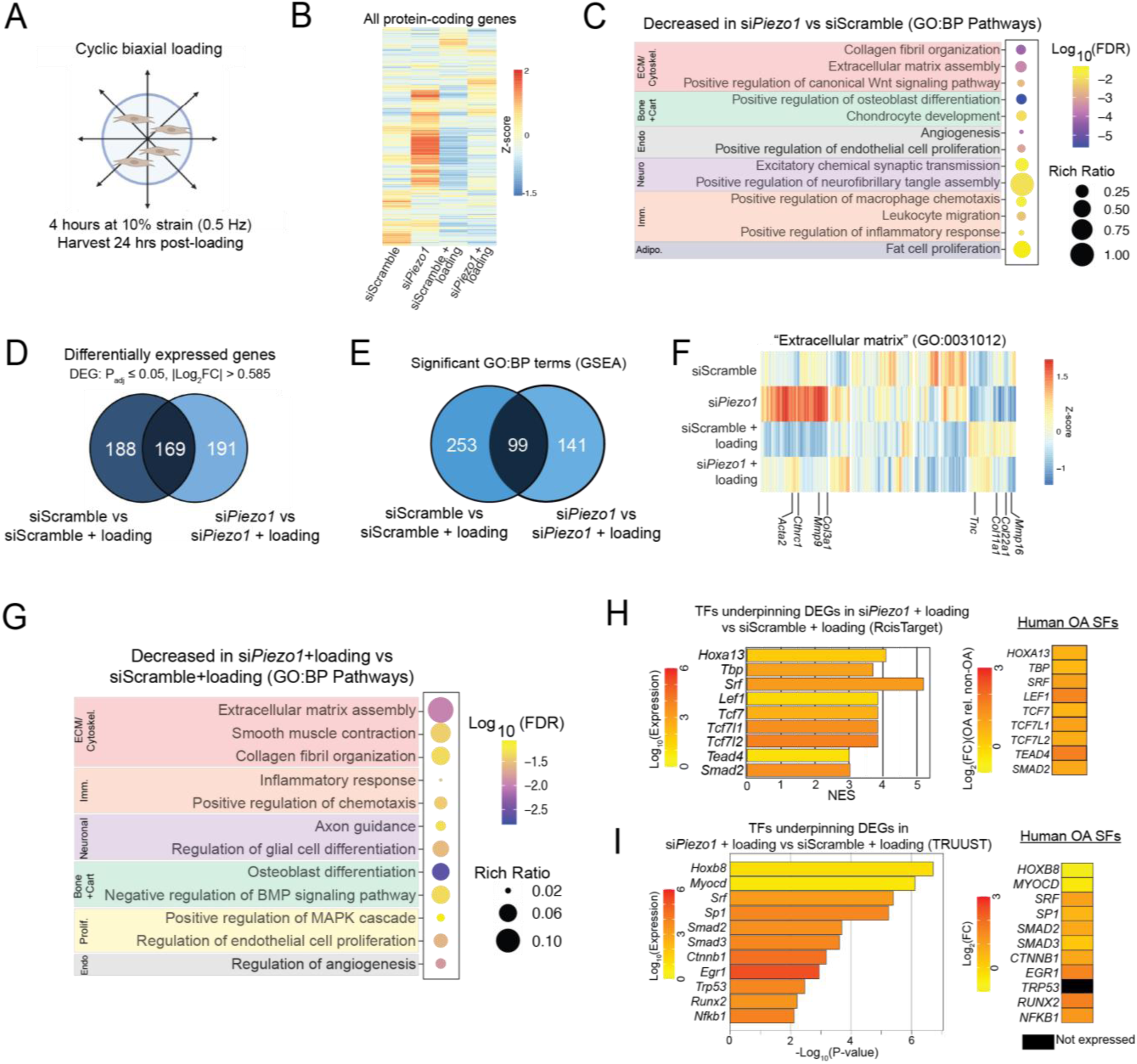
PIEZO1 regulates the mechanical loading response of SFs. **(A)** Schematic of cyclic biaxial tensile loading regimen conducted on murine SFs. Bulk RNA-seq was conducted on n=4 biological replicates (unique isolations) per condition. **(B)** Gene expression by RNA-seq of all protein-coding genes across experimental conditions. Aggregate relative expression of n=4 biological replicates per condition is plotted. **(C)** Gene Ontology:Biological Processes (GO:BP) terms (p_adj_<0.05) enriched in DEGs downregulated in si*Piezo1* versus siScramble groups. Statistical overrepresentation of significant downregulated DEGs defined by p_adj_<0.05, |Log_2_FC|>0.585. **(D)** Venn diagram illustrating overlap of loading-induced DEGs in SFs with and without concomitant treatment with si*Piezo1*. **(E)** Venn diagram illustrating overlap of significant loading-induced GO:BP pathways (p_adj_<0.05, via GSEA) in SFs with and without concomitant treatment with si*Piezo1.* **(F)** Relative expression of all genes comprising the “Extracellular matrix” GO:BP term (GO:0031012) across experimental conditions. Aggregate relative expression of n=4 biological replicates per condition is plotted. **(G)** Selected GO:BP terms (p_adj_<0.05) enriched in DEGs downregulated in si*Piezo1-*treated loaded SFs versus siScramble-treated loaded SFs. Statistical overrepresentation of significant downregulated DEGs defined by p_adj_<0.05, |Log_2_FC|>0.585. **(H)** Left: Normalized enrichment scores (NES) of high-quality, direct annotation-derived TFs (NES>3) predicted to regulate DEGs in si*Piezo1-*treated loaded SFs versus siScramble-treated loaded SFs, derived via RCisTarget. Right: Differential expression of corresponding human genes, plotted as Log_2_FC in OA vs non-OA human SFs. **I)** Left: Pval of TFs predicted to regulate DEGs in si*Piezo1-*treated loaded SFs versus siScramble-treated loaded SFs, derived via TRUUST. Right: Differential expression of corresponding human genes, plotted as log_2_ fold-change in OA vs non-OA human synovial fibroblasts.

Of the DEGs induced by loading (357 in siScramble-treated SFs and 360 in si*Piezo1*-treated SFs), less than half (169) were shared, indicating a substantial role of PIEZO1 in mediating the SF loading response (**Fig 4D**). Among loading-induced genes, those most downregulated in loaded si*Piezo1* versus loaded siScramble SFs were associated with fibrosis and ECM production (*Tnmd, Col14a1, Bglap2, Bmp4*), myofibroblast markers (*Cnn1, Lmod1*), inflammation (*Cxcl1*), and cell cycle regulation (*Igfbp2, Ccno*) (**Fig S5D, Supplemental Table 9**). Gene-set enrichment analysis (GSEA) demonstrated large *Piezo1* silencing-induced perturbations of the pathways activated by loading (**Fig 4E**), including terms relevant to fibrosis and ECM remodeling such as the pathway “Extracellular Matrix” (**Fig 4F**). Strikingly, several myofibroblast and fibrosis markers in this pathway (*Acta2*, *Cthrc1*, *Mmp9*, *Col3a1*) and cell adhesion-related genes such as *Itga3* were paradoxically increased in si*Piezo1*. Given the clear pro-myofibroblastic effect of PIEZO1 agonism in SFs (**Figure 3**), this may be a compensatory response to *Piezo1* silencing, reflecting diminished ability of SFs to mechanosense their environment. Pathways related to ECM and collagen production, calcium ion transport, cellular contraction, neurotrophic signaling, osteoblastic differentiation, angiogenesis, and inflammatory activity mapped to downregulated DEGs in loaded si*Piezo1*-versus siScramble-treated SFs (**Fig 4G**).

Inferring molecular regulation of these effects, RcisTarget and TRUUST analyses both predicted multiple Wnt-related (e.g. TCF7, LEF1, CTNNB1), YAP1-related and TGF-β-related (e.g. TEAD4, SMAD2/3) TFs in mediating the PIEZO1 response to mechanical loading, as their binding motifs were overrepresented in genes differentially expressed in si*Piezo1* vs siScramble loaded SFs (**Fig 4H**, **Fig 4I**). Several genes encoding these TFs were also differentially expressed in human OA versus non-OA SFs, suggesting differential PIEZO1-related signaling in human OA tissue. Taken together, these data support the involvement of PIEZO1 in the signal transduction of mechanical loading in SFs.

### PIEZO1 overactivation induces a pro-fibrotic and anti-adipogenic synovial response in vivo

We next assessed whether acute intra-articular PIEZO1 activation alone is sufficient to induce a pathological synovial phenotype in *in vivo*. Mechanical activation of PIEZO1 in PTOA synovium may occur acutely (at the time of injury) or chronically (due to ECM stiffness changes during disease progression). To simulate acute PIEZO1 overactivation pharmacologically, naïve male and female C57Bl/6 mice were administered 5 consecutive days of intra-articular Yoda1 (4 mg/kg) or vehicle (5% DMSO) injections (**Fig 5A**). This acute PIEZO1 activation regimen was sufficient to induce long-term synovial remodeling, as histological scoring at 28d after the first injection demonstrated significantly increased synovitis scores in both male and female Yoda1-treated mice (**Fig 5B-E, Fig S6A, B**). This synovitis was strongly driven by increased lining layer hyperplasia and more abundant connective tissue in the sublining and IFP compartments in Yoda1-compared to vehicle-injected joints (**Fig 5B-C**). Histologically, we did not observe any significant Yoda1-induced effects on articular cartilage, meniscus, or bone (**Fig S6A**).

**Fig 5.**
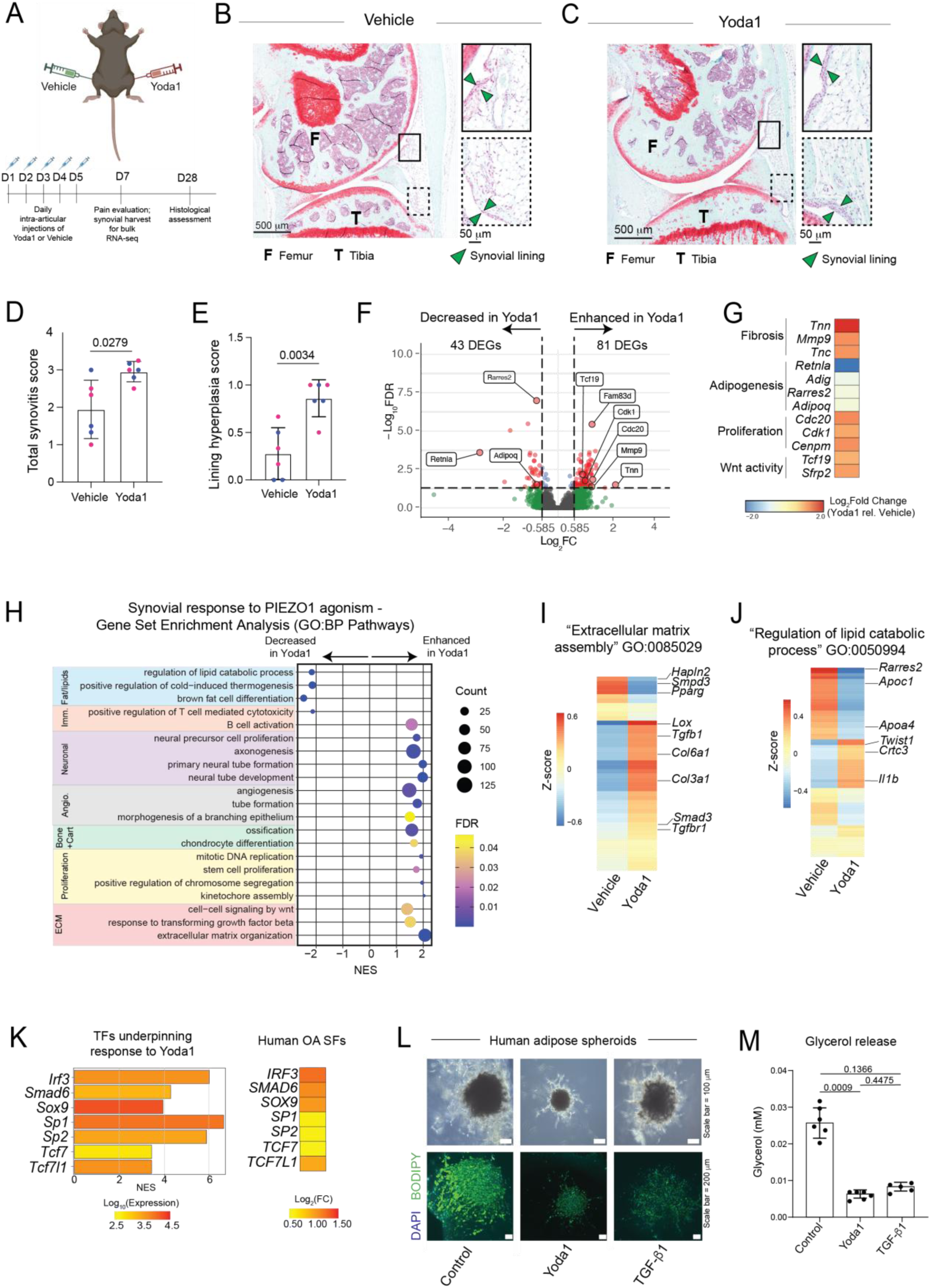
PIEZO1 overactivation induces a pro-fibrotic and anti-adipogenic synovial response in vivo. **(A)** Schematic illustrating experimental design and timeline of intra-articular vehicle or Yoda1 injections. **(B-C)** Representative sagittal Safranin-O/Fast Green-stained sections of the medial joint compartment in Vehicle (**B**) and Yoda1 (**C**)-injected joints, collected at 28 days following the first injection. Zoom insets illustrate anterior synovium and IFP, with the lining marked by arrow heads. **(D-E)** Total histological synovitis score (D) and histological lining hyperplasia subscore (E) in vehicle and Yoda1-injected joints. n=6 per condition (3 male, 3 female). Two-tailed paired t-test. **F)** Volcano plot from bulk RNA-seq of synovium from vehicle vs Yoda1-injected joints, collected at 7 days following the first injection. DEGs are colored in red and defined as p_adj_ < 0.05, |Log_2_FC| > 0.585. Bulk RNA-seq was conducted on matched bilateral synovia from n=6 mice (3 male, 3 female). **G)** Differential expression in log_2_ fold-change of selected DEGs upregulated in synovium from Yoda1-injected vs vehicle-injected joints. **(H)** Selected GO:BP terms (p_adj_ < 0.05) differentially enriched in synovium from Yoda1-injected vs vehicle-injected joints. Gene Set Enrichment Analysis (GSEA). n=6 per condition (3 male, 3 female). **(I-J)** Relative expression of all genes comprising the GO:BP “Extracellular matrix assembly” (I) and “Regulation of lipid catabolic process” (J) pathway terms. Aggregate relative expression of n=6 per condition (3 male, 3 female) is plotted. **(K)** Normalized enrichment scores (NES) of high-quality, direct annotation-derived TFs (NES > 3) predicted to regulate DEGs in Yoda1-injected vs vehicle-injected synovia, derived via RCisTarget. Right: Differential expression of corresponding human genes, plotted as log_2_ fold-change in OA vs non-OA human synovial fibroblasts. **(L)** Brightfield and fluorescent microscopy of human adipose spheroids. DPP4+ preadipocytes were formed into spheroids and exposed to adipogenic medium for 14 days, with or without supplementation of 10 µM Yoda1 or 10 ng/mL TGF-β1. Lipid is labeled by BODIPY and nuclei by DAPI staining. **(M)** Glycerol concentration in conditioned medium at day 14 from spheroids in (L). n=6 spheroids per condition. One-way ANOVA with Tukey’s post-hoc testing.

We then employed bulk RNA-seq to assess the molecular programs underpinning this remodeling. Whole synovia, including the IFP, were homogenized for RNA isolation and RNA-seq at d7 (48h after the final injection). Given the neurotrophic-related pathways in our *in vitro* findings, we performed blinded knee hyperalgesia assessments at d7, but Yoda1 injections did not alter knee withdrawal thresholds (**Fig S7A).** After quality control **(Supplemental Table 10),** synovia from Yoda1-injected limbs exhibited 81 upregulated and 43 downregulated DEGs (Padj<0.05, |Log2FC|>0.585, FKPM>1) versus vehicle-injected joints (**Fig 5F, Fig S7B, Supplemental Tables 11**). Notably, upregulated DEGs included cWnt-related genes of the *Tcf*-family, proliferation-related genes (*Cdk1*, *Cdc20*), and pro-fibrotic genes (*Tnc*, *Tnn*, *Sfrp2*) (**Fig 5F-G**). Downregulated DEGs had a marked adipose signature, with adipogenic genes (*Rarres2*, *Retnla*, *Adipoq*) reduced in synovia of Yoda1-treated mice. GSEA demonstrated enriched pathways related to the ECM, TGF-β and Wnt signaling, cellular proliferation, bone and cartilage formation, angiogenesis, and neurotrophic signaling in Yoda1-treated synovia (**Fig 5H**). Downregulated GSEA terms related to adipose and lipid functions, indicating an anti-adipogenic effect of PIEZO1 agonism reminiscent of the IFP adipose attrition we observe in the ACLR model^39^. Statistical overrepresentation analysis also identified enriched pathways related to cell proliferation and downregulated pathways related to adipose tissue and lipid metabolism (**Fig S7C-D**). Significant leading-edge genes in the Yoda1-enhanced “Extracellular matrix assembly” pathway showed Yoda1-induced upregulation of fibrosis- and TGFβ-relevant genes (*Tgfb1*, *Tgfbr1*, *Smad3 Col3a1*, *Lox*), with concurrent downregulation of the adipogenic markers *Pparg* and *Smpd3* (**Fig 5I**). Concordantly, downregulated genes in the Yoda1-decreased “Regulation of lipid catabolic process” pathway included the apolipoproteins *Apoa4*, *Apoc1* and *Rarres2*, which encodes the adipokine chemerin, with upregulation of the cytokine gene *Il1b* and the anti-adipogenic *Twist1* (**Fig 5J**).

Using reference scRNA-seq data from murine synovium, leading edge genes of significant GSEA pathways mapped across multiple synovial cell types, suggesting synovial PIEZO1 agonism directly or indirectly affects multiple cellular subsets. Fibroblasts contributed most strongly to the bone and cartilage, ECM and cytoskeleton, and angiogenesis-related modules activated by Yoda1 injection **(Fig S7E**). Transcription factor binding motif analysis of Yoda1-induced genes identified SMAD6, the SP- and TCF-families, the chondrogenesis and fibrosis-relevant TF SOX9, and the interferon regulator IRF3 as putative mediators of the Yoda1 response (**Fig 5K**).

Lastly, given the strong anti-adipogenic signature in Yoda1-injected joints and the progressive fibrosis of intra-articular adipose tissue in murine and human PTOA and other chronic fibrotic diseases, we tested whether PIEZO1 agonism directly blunts adipogenesis of stromal progenitors. We generated adipose spheroids from human DPP4+ preadipocytes to model the *Dpp4*+ *Pi16+* synovial progenitors underpinning myofibroblast-like SF emergence after ACLR. While control spheroids in adipogenic medium progressively increased in size and formed robust lipid droplets after 14 days, supplementation with 10 µM Yoda1 markedly inhibited spheroid growth and lipid accumulation (**Fig. 5L, Fig. S8A**). These effects exceeded those of 10 ng/mL TGF-β1, a positive control for pro-fibrogenic and anti-adipogenic induction. To corroborate the downregulation of lipid catabolism terms in Yoda1-injected joints, we measured glycerol release as a measure of functional lipolysis. Yoda1-supplemented spheroids exhibited an ∼80% reduction in glycerol release, comparable to TGF-β1-treated spheroids (**Fig. 5M**). These results confirm a strong, direct anti-adipogenic effect of PIEZO1 agonism in DPP4+ stromal progenitors.

Taken together, PIEZO1 overactivation initiates pro-fibrotic, pro-proliferative and anti-adipogenic signaling that drives synovitis-relevant tissue remodeling in vivo.

## Discussion

Based on synovial pathology’s strong association with patient-reported pain and cartilage loss and disease risk in OA^60–63^, there is a growing focus on identifying the mechanisms that drive chronic OA synovitis. Here we demonstrated the emergence of YAP1-active myofibroblast-like SFs after injury and showed that PIEZO1 is a critical mechanoreceptor activating myofibroblastic differentiation, mediates pathological mechanical loading responses, and can drive a pro-fibrotic, pro-inflammatory, and anti-adipogenic response *in vivo*.

Prior scRNA-seq studies focused on disease-associated SFs in joint diseases including OA and RA^8,11,38,64^ have not resolved their anatomic distribution. Using Xenium, Bhamidipati et al. demonstrated enrichment of COMP-Hi fibrogenic SFs in RA treatment non-responders, spatially mapping these cells to a perivascular compartment^65^. Single-cell spatial mapping of PTOA synovium, especially mapping of the early emergence of disease-associated cellular subsets, had not been conducted. Using Xenium 5K spatial transcriptomics, we identified anatomic niches enriched with myofibroblast-like SFs in PTOA, primarily dense fibrotic tissue and perivascular regions in the sublining and IFP, similar to the observations of Bhamidipati et al^65^. This suggests that interactions between endothelial cells and SFs may govern myofibroblastic activation, as shown in RA^64–66^. While some myofibroblast-like SFs were observed in the immediate sublining, we did not observe them to strongly populate the lining. Spatially dependent cell-cell communication involving these myofibroblast-like SFs should be described in future studies. Consistent with observations in other disease contexts^67–70^, our spatial data also revealed that these cells exhibit a pro-inflammatory and neurotrophic transcriptional signature, besides their overactive ECM- and cytoskeletal contraction-related programs. This was independently corroborated in murine and human SF scRNA-seq (e.g. overrepresented “Positive regulation of leukocyte chemotaxis” and “Axon guidance” GO:BP pathways in human myofibroblast-like SFs). Biologically, these findings suggest that myofibroblast-like SFs may also be perpetuating low-level synovial inflammation and participating in pain-relevant neurotrophic signaling in OA.

Stromal progenitors and the mechanisms driving their fibrogenic activation are of significant interest to the development of anti-fibrotic therapies. Prior work showed that *Dpp4+ Pi16+* synovial progenitors transition to multiple pathological SF subsets, including myofibroblast-like SFs^8,15^. A similar fate commitment by universal fibroblasts has been demonstrated in other fibrotic conditions such as in the heart and lungs^71,72^. Leveraging trajectory analysis, we identified that the fate commitment of activated *Dpp4+ Pi16+* progenitors towards a myofibroblastic phenotype involves activation of a YAP1-associated regulon. This suggested the involvement of upstream mechanoreceptors governing YAP1. While YAP1 has been described as a mechanosensitive transcriptional mediator in joint injury- and inflammation-induced synovial hyperplasia^34,35,37^, it does not directly sense mechanical signals, and the specific upstream mechanoreceptors remain unclear. Our cross-species OA SF atlas demonstrated that *Piezo1/PIEZO1* is the most highly expressed and enriched mechanoreceptor in murine PTOA and human OA SFs compared to controls. *Piezo1* was upregulated along the trajectory in murine synovium, and *Piezo1* and *Yap1* expressions were highly correlated across murine SF subsets. Connecting these factors functionally, we confirmed that both PIEZO1 agonism and culture substrate stiffness increased YAP1 nuclear content, and that PIEZO1 agonism drove myofibroblastic activation in a YAP1-dependent manner. These observations suggest that myofibroblastic activation of synovial progenitors is driven, at least in part, by overactive SF mechanosensing, potentially due to increased ECM stiffness as measured by AFM indentation. Establishing the causality between ECM stiffness changes, enhanced mechanosensing, and myofibroblastic activation is an important next step. As various inflammatory factors may also prime quiescent fibroblasts towards myofibroblastic activation^73^, the robust joint injury-induced inflammatory response likely also contributes to this sequence. In this context, inflammatory priming by IL-1α was shown to sensitize articular chondrocytes to PIEZO1-mediated mechanosensing, underpinning the pathological chondrocyte phenotype in OA^74^.

To characterize the role of PIEZO1 in the SF’s mechanosensing of loading, relevant to the dynamic mechanical nature of synovial joints and the overloading that occurs during PTOA progression, we applied cyclic biaxial tension (10% strain at 0.5 Hz) to murine SFs with and without intact *Piezo1*. siRNA-mediated *Piezo1* silencing had a strong effect on both unloaded and loaded cells, and *Piezo1*-silenced SFs shared less than half of loading-induced DEGs compared to control SFs. These findings are consistent with the canonical role of PIEZO1 as a mechanosensitive ion channel that activates under membrane deformation, and the large effect of *Piezo1* silencing is likely due to it being highly expressed in SFs, as our human and murine scRNA-seq findings demonstrated. In both unloaded and loaded cells, *Piezo1* silencing was associated with downregulated pathways related to ECM organization and collagen production, osteochondral differentiation, inflammatory signaling, and angiogenesis, linking *Piezo1* to key SF phenotypes relevant to fibrosis and inflammation in PTOA. In loaded cells, *Piezo1* silencing downregulated proliferation-related genes and pathways, potentially via its intersection with cWnt signaling. In skin wound healing, PIEZO1 was shown to control proliferation of keratinocytes^75^, and in growth plate chondrocytes, PIEZO1 mediated proliferation through a NFAT-YAP1-β-catenin axis^76^. In the context of bladder outlet obstruction, PIEZO1 mediated fibroblast proliferation and migration via YAP1/GLS1^77^. Inferred transcription factors associated with genes differentially expressed due to *Piezo1* silencing included those involved in cWnt and TGFβ-SMAD signaling, including genes of YAP1 co-factors such as *Tead4*, further connecting PIEZO1 activity to these fibrosis-relevant pathways. Unexpectedly, we also observed that *Piezo1* silencing induced upregulation of ECM-, cytoskeletal contraction-, and cell adhesion-related genes, including upregulated *Acta2, Col3a1, Cthrc1,* and *Itga3*, which we suspect may be a compensatory response due to a decreased capacity of SFs to mechanosense their environment via PIEZO1. PIEZO1-dependent mechanosensing has been found to be critical in promotion of fibrosis in tissues such as the kidneys, lungs, and skin^54,78,79^. While our studies focused on dynamic tensile loading, the mechanical loading observed by SFs *in vivo* is more complex and dependent upon various factors such as anatomic location and tissue disease status. Thus, future studies should investigate how PIEZO1 governs the SF response to other mechanical perturbations.

Acute intra-articular PIEZO1 overactivation via a five-day regimen of Yoda1 joint injections induced synovitis in healthy murine joints, evidenced by increased histological synovitis scores and a pro-fibrotic, pro-proliferative, and anti-adipogenic synovial transcriptomic response. This suggests that overactive PIEZO1-mediated mechanosensing is sufficient by itself to induce a pathological synovial response *in vivo*. We used Yoda1 injections to mimic overactive PIEZO1 mechanosensing in the acute post-injury period. This was informed by our present and published findings demonstrating a robust early fibroblastic injury response in synovium, including enrichment of pathways related to mechanosensing such as “response to fluid shear stress”, “actin crosslink formation”, and “cellular response to mechanical stimulus”. Our findings are consistent with an acute response resulting in long-term remodeling, such as the acute activation of proliferation genes and pathways corresponding to synovial lining hyperplasia observed in Yoda1-injected joints. In addition to upregulated genes and pathways associated with fibrosis, proliferation, and cWnt activity, including myofibroblast marker genes, we also observed a robust anti-adipogenic transcriptomic response to Yoda1. We functionally validated this observation by demonstrating that Yoda1 supplementation of adipogenic medium robustly blocks adipogenesis and glycerol production of DPP4+ human adipose spheroids. No prior studies have described the role of PIEZO1 in the regulation of intra-articular adipose tissue, but anti-adipogenic effects of PIEZO1 activation have been demonstrated in bone marrow and perivascular adipose tissue^80,81^. However, PIEZO1 activation by mature adipocytes was also found to enhance obesogenic adipogenesis of preadipocytes^82^, suggesting a context-dependent role of PIEZO1 mechanosensing in adipose regulation. Our findings are relevant to the attrition of intra-articular adipose tissue and downregulation of adipose-related pathways we observed in the murine ACLR model^39,40^. Although histological scoring did not demonstrate massive loss of intra-articular adipose tissue, we observed increased connective tissue deposition in the synovial sublining and IFP compartments. These findings and the anti-adipogenic effects of PIEZO1 agonism in adipose spheroids *in vitro* are consistent with fibrogenic activation of synovial progenitors, but the direct effects of PIEZO1 agonism on mature synovial adipocytes should be assessed in future studies. Furthermore, while RNAseq also demonstrated a Yoda1-induced neurotrophic response, e.g. upregulated “axonogenesis” GO:BP pathway, we did not observe increased knee hyperalgesia two days after the last injection. Thus, longer-term sensitization assessments should be performed to assess whether PIEZO1 overactivation manifests in a delayed hyperalgesia response. A limitation of this experiment was the inability to specifically target synovium, and cell type-specific modulation of PIEZO1 signaling should be conducted in future studies to pinpoint the effects of PIEZO1 mechanosensing to specific cells.

In summary, this study identified the activation of a PIEZO1-YAP1 signaling axis in the emergence of fibrogenic, myofibroblast-like synovial fibroblasts after joint injury and demonstrated that PIEZO1 activation regulates myofibroblastic activation and mechanosensing of SFs. Given the relevance of synovitis in the clinic, this provides a potentially targetable mechanism for PTOA treatment.

## Methods

### Mice

Mice were housed communally in ventilated cages and provided *ad libitum* access to chow and water in a 12h light-dark cycle. All mice were between 12 – 16 weeks of age at the time of experimentation. C57Bl/6J mice were sourced from Jackson Laboratories (Strain #000664). *Acta2-*Cre^ERT2^ mice were kindly provided by Dr. Ivo Kalajzic^83^ (Strain #032758) and crossed with Ai9 mice (Strain #007909) to generate a tamoxifen-inducible tdTomato-expressing reporter of *Acta2*-expressing cells, henceforth termed *Acta2*-Cre^ERT2^;tdTom. Mice were euthanized via CO_2_ asphyxiation or by exsanguination under deep isoflurane-induced anesthesia during terminal cardiac perfusions. All procedures involving mice were conducted under approval from the Institutional Animal Care and Use Committee (IACUC) of the University of Michigan (#PRO00011634).

### Anterior cruciate ligament rupture model of post-traumatic osteoarthritis

Joint injury was administered to mice via a noninvasive, tibial compression-based anterior cruciate ligament rupture (ACLR) model, as previously described^40,84,85^. Briefly, following induction of inhaled isoflurane anesthesia, the flexed hindlimb was positioned in a custom fixture and subjected to preloading, preconditioning, and a rapid 1.5 mm compressive displacement to induce tibiofemoral subluxation and complete ACL^86,87^. Following injury, mice are administered a single subcutaneous dose of carprofen (5 mg/kg). ACLR procedures were performed using either a modified CellScale Univert S2 or an Electroforce 3300AT mechanical testing system, which have been shown to have high concordance^85^.

### Structural Histology

Whole hindlimbs were fixed and then decalcified in 7.4 pH Na^2+^ EDTA for 14d, followed by standard paraffin processing and embedding. Sagittal sections (5 µm thickness) spanning the knee joint were obtained and stained with either Safranin-O/Fast-Green or Masson’s Trichrome^88^. Six to nine sections were stained and analyzed per joint. Stained sections were imaged at 20x using either a Keyence BZ-X810 or a Nikon Eclipse Ni E800 with DS-Ri2 camera.

### Histological Scoring

Microscopic images of sagittal Safranin-O/Fast-Green-stained sections spanning the medial condyle were scored for PTOA and synovitis severity in a blinded fashion, as previously described^89^. Scoring categories and weights are listed in Supplemental Table 12. Scores were averaged across 5-6 sections per joint.

### Atomic force microscopy (AFM)-based nanoindentation

AFM-based nanoindentation was performed to quantify stiffness of the synovial ECM. Following euthanasia, whole hindlimbs were harvested, trimmed of excess muscle, and bathed in a sucrose solution overnight. Limbs were then flash frozen in OCT. Following embedding, 40 µm-thick sagittal cryosections (1 section per limb) were taken from the medial compartment of Sham and ACLR 28d male and female mice (n=10 mice per condition) using the Kawamoto’s film-stabilization method^90^. On each section, 15-20 indentations were performed in the sublining synovium of the anterior compartment, spanning the entire cranial-caudal length of the tissue to account for spatial heterogeneity in ECM stiffness. Polystyrene microspherical tips (R ≈ 12.45 μm, nominal k ≈ 0.6 N/m, HQ:NSC36/tipless/Cr-Au, cantilever C, NanoAndMore) were utilized for nanoindentation using a Dimension Icon AFM (Bruker Nano). Indentation was applied at a rate of 10 µm/s with a maximum load of ≈120 nN. Effective indentation modulus (E_ind_) was computed using the finite thickness-corrected Hertz model (Poisson’s ratio (ν) assumed to be 0.4), all as previously described^11^.

### Flow cytometry of Acta2-Cre^ERT2^;tdTom reporter mice

*Acta2/*αSMA-expressing synovial fibroblasts were quantified in synovia from ACLR 7d and respective contralateral joints using flow cytometry. *Acta2-*Cre^ERT2^;tdTom mice (n = 4 males and 4 females) underwent ACLR and received intraperitoneal tamoxifen injections (75 mg/kg) on days −2, 0, and +2 post-ACLR. Following euthanasia, whole synovium, inclusive of Hoffa’s fat pad, was microdissected and enzymatically digested in DMEM containing 400 µg/mL DNaseI (Sigma DN25), 400 µg/mL Liberase TM (Sigma 5401127001), and 400 µg/mL Collagenase IV (Sigma C5138) for 30 min at 37°C. During digestion, samples were vortexed every 15 minutes. Digestates were then filtered (40 mm mesh size) and centrifuged at 500 x g for 5 mins to yield a single-cell suspension, as previously described^8,84^. To assess cellular viability, cells were stained with TO-PRO-3 (Invitrogen). Cold FACS buffer (PBS, 2% FBS, 1 µM EDTA) was used throughout staining and during cytometric analyses. Non-specific antibody binding was blocked using TruStain FcX (Biolegend). CD31-PECy7 (Biolegend) was used to label endothelial cells, and CD45-BV650 (Biolegend) was used to label hematopoietic cells. Thy1-PEDaz594 (Biolegend) was used to mark fibroblasts, and CD146-BV605 (Biolegend) was used to identify and exclude αSMA-expressing pericytes. Cells with negative expression of CD31, CD45, and CD146 but positive expression of Thy1 and tdTom were identified as myofibroblast-like SFs. Unstained, single-stained, and fluorescence-minus-one (FMO) controls from WT synovia were included alongside samples to assist with gating. Compensation and acquisition were performed using a BD LSRFortessa cytometer with FACSDiva software (BD Biosciences). Data analysis was performed using FlowJo v10 (TreeStar/BD Biosciences).

### Isolation and culture of primary synovial fibroblasts

For *in vitro* experiments, primary SFs were isolated from murine hindpaw synovium, as previously described^8^. Briefly, following euthanasia, hindpaws were removed just above the ankle joint, and skin and nails were removed. Longitudinal cuts were made using a scalpel on the ventral and dorsal sides of the paws to separate the hindpaw joints. Hindpaws were then enzymatically digested at 37°C in DMEM containing 400 mg/mL DNAseI (Sigma DN25), 400 mg/mL Liberase (Sigma 5401127001), and 400 mg/mL Collagenase IV (Sigma C5138) in DMEM). Samples were vortexed every 5 min for 45 min, then digestates were strained (70 mm mesh size) and centrifuged at 500 x g for 5 mins to yield single-cell suspensions. Cells were plated and cultured in complete media (10% FBS, 1% P/S, 0.2% Amphotecerin/Fungizone in DMEM) up to P3. Prior to experiments, SFs were cultured in serum-reduced (1% FBS) medium overnight. For experimental media, 0% FBS (ICC, SuperTopFlash) or 1% FBS (wound healing assay, gel contraction assay) media was used.

### Small interfering RNA (siRNA) transfection

Primary SFs at P2 were seeded into T75 flasks (∼35,000 SFs/cm^2^) and raised to ∼80% confluency prior to transfection with 10 nM si*Piezo1* (pooled siRNA, siGenome, M-061455-01-0005) or non-targeting siScramble (ON-TARGETplus Non-targeting Control Pool siRNA, Horizon Discovery D-001810-10-05) overnight with the DharmaFECT 1 Transfection Reagent (Horizon Discovery T-2001-03) in OPTI-MEM. Media was replaced with complete media and cells were passaged to p3 for experiments. Knockdown was confirmed by qPCR prior to experimentation.

### HEK293 SuperTopFlash Assay

SuperTopFlash (STF) HEK293 cells (Kerafast, EJH014) cells were cultured in DMEM supplemented with (10% FBS, 1% P/S, 0.2% Amphotecerin/Fungizone, and 4 mL/mL G418 Sulfate Antibiotic Solution (Mirus MIR5920). Experiments were performed at p6-8 at a cell density of ∼60,000 cells/cm^2^. Cells were treated combinatorially with Yoda1 (10 µM), or the selective YAP1 inhibitor Verteporfin (0.5 µM) for 1d prior to performing the assay. OneGLO lysis buffer, 1X (Promega E2661) was added to HEK293 experimental samples, which were shaken orbitally at 100 RPM for 3 minutes to facilitate lysis. ONE-Glo Luciferase Assay Buffer (Promega E605A) was then added to each sample, and luminescence was read using a Synergy HTX Multi-Mode Microplate Reader (Agilent BioTek). Luminescence reads were normalized to viability based on Alamar Blue fluorescence.

### In vitro Fibroblast Phenotyping

#### Gene expression analysis by quantitative real-time PCR (qPCR)

Cells were lysed and homogenized in TRIzol. RNA isolation was performed by phenol chloroform extraction. cDNA synthesis was performed using the random primers method and a High-Capacity cDNA Reverse Transcription Kit (Applied Biosystems, CA, USA), and cDNA was combined with SYBR Green Master Mix (Applied Biosystems, CA, USA) for qPCR analysis on a Bio-Rad T100 Thermal Cycler (Bio-Rad, Berkeley, CA, USA). Ct values were normalized using the 2^−ΔΔCT^ method, using *Atp5b* as the housekeeper gene. Primer sequences are listed in Supplemental Table 13.

#### Immunocytochemistry

Primary SFs were cultured to ∼80% confluency on tissue culture plastic or Ibidi 35 mm high ESS micro-dishes (1.5 kPa) (Ibidi 81291) coated with fibrin (Sigma Aldrich F8630) and fixed with 4% paraformaldehyde for 15 mins. To coat Ibidi micro-dishes, fibrinogen was dissolved to 0.8 mg/mL in 1M Tris-HCl. 0.8 mL of the resulting fibrin solution was added to each micro-dish at 37°C for 2 hours. Dishes were then washed in triplicate with PBS before SFs were plated. For immunostaining of αSMA and F-actin, cells were then permeabilized with 0.2% Triton X-100 solution for 12 mins (Sigma Aldrich T8787) and blocked for 2h in a solution containing 10% w/v bovine serum albumin (Fisher BP9706), 5% v/v normal goat serum (Fisher 16210064), and 2% w/v glycine (Fisher G48-212). Conjugated αSMA antibody (eBiosciences 53-9760-80) and Phalloidin-iFluor 750 reagent (Abcam ab176762) each diluted 1:500 in blocking solution were added to cells overnight at 4°C. For immunostaining of YAP1 and β-catenin, cells were fixed with 4% paraformaldehyde for 15 mins, then permeabilized with 0.2% Triton X-100 solution for 10 mins (Sigma Aldrich T8787) and blocked for 1hr in a solution containing 10% w/v bovine serum albumin. Primary anti-YAP1 antibody (Santa Cruz sc-101199) was diluted 1:400 in PBS containing 1% w/v bovine serum albumin and added to cells overnight at 4°C. Primary antibody specific to non-phosphorylated (active) β-catenin (Cell Signaling 8814S) was diluted 1:400 PBS containing 1% w/v bovine serum albumin added to cells overnight at 4°C. Secondary antibody (Cell Signaling A21200) diluted to 1:500 in 10% BSA was then added for 2h at room temperature. Cells were then counterstained with Hoechst nuclear stain (Invitrogen 33342) and images were acquired using a Lionheart FX (Biotek) microscope. To assess total cellular αSMA and F-actin protein, average fluorescent signal intensity was quantified by thresholding the entire cell using the Otsu method via MATLAB R2022a (Mathworks). To assess nuclear YAP1 and β-catenin protein, binary nuclear masks were derived from Hoechst-stained images via the Otsu method, and average fluorescent signal intensity of YAP1 or β-catenin were quantified within nuclear masks.

#### Scratch wound assay

Primary SFs were cultured on 24 well plates up to ∼90% confluency. A scratch was introduced down the centerline of each well using a p200 pipette tip. SFs were gently rinsed with PBS and then given experimental treatments. Fibroblast phenotyping involved combinatorial treatments with Yoda1 and Verteporfin, or si*Piezo1* and TGF-β1. SFs derived from healthy murine hindpaws were treated with either vehicle (DMSO), Yoda1 (10 µM), the selective YAP1 inhibitor Verteporfin (0.5 µM), or a combination of Yoda1 and Verteporfin for 1d prior to introducing the scratch. Treatment with si*Piezo1* or non-targeting siScramble was performed overnight at p2 prior to passaging and plating at ∼80% confluency. SFs were then treated with either vehicle or TGF-β1 (10 ng/mL) for 1d prior to introducing the scratch. Scratch sites were imaged longitudinally at 0, 4, 12, and 24 hr post-scratch, and the proportion of the recovered wound site area was quantified with ImageJ. The area under the curve (AUC) of the proportion of recovered wound versus time was calculated and compared between conditions.

#### Collagen gel contraction assay

Primary SFs at P2 were cultured on T75 plates to ∼90% confluency, trypsinized, and seeded into a collagen gel (Cell Biolabs, Inc., CBA-021) at ∼1×10^6^ SF/mL, according to manufacturer’s instructions. Collagen-embedded SFs were maintained in complete media for 24h, serum starved overnight (1% FBS), and exposed to experimental treatments. As described above, SFs were treated combinatorially with Yoda1 and Verteporfin, or si*Piezo1* and TGF-β1. For transfection experiments, treatment with siScramble or si*Piezo1* occurred overnight prior to trypsinization at P2. Treatment with Yoda1, Verteporfin, and TGF-β1 and respective vehicles was applied one day after embedding SFs into gels. Following 24h of exposure to experimental conditions, gels were separated from the wells using a spatula and media was replaced with starvation media. Images were then acquired longitudinally (0, 4, 8, 12, and 24h after release) and gel surface areas were quantified using ImageJ. The proportion of original surface area at t=0 was calculated at each timepoint. The AUC of the proportion of original surface area versus time was calculated and compared between conditions.

### *In vitro* mechanical loading of SFs

Primary SFs at P3 were seeded onto CellSoft BioFlex plates (10 kPa stiffness) coated with collagen I (FlexCell International). As described above, si*Piezo1* silencing or non-targeting siScramble treatment was conducted overnight at p2 prior to seeding onto BioFlex plates. Wells were seeded at ∼35,000 SFs/cm^2^ At ∼80% confluency, SFs were subjected to cyclical biaxial loading (10% elongation, 0.5 Hz) for 4h using the Flexcell FX4000T system on CellSoftTM 6 well BioFlex® plates (10 kPa) coated with Collagen I. SFs were then evaluated for viability using the Alamar Blue assay and homogenized in TRIzol 24h after the end of the loading bout, then submitted for bulk RNA-sequencing.

### Intra-articular Yoda1 injections

To overactivate PIEZO1 signaling *in vivo,* Yoda1 was injected intra-articularly for five consecutive days. Naïve male and female C57Bl/6J mice (n=6, 3 male and 3 female) underwent isoflurane-induced anesthesia and received five consecutive daily injections of either Yoda1 (4 mg/kg at 10 µM in 5% DMSO) in one joint and vehicle (equivalent volume of 5% DMSO) in the contralateral joint. Injections were administered using a 33G needle and a 10 mL microsyringe (Hamilton). Limb designations were randomly assigned. All injections and assessments were conducted in a blinded fashion. At 7 or 28 days following the first injection, mice were euthanized by CO_2_ asphyxiation. Whole synovium samples, inclusive of Hoffa’s fat pad, were collected in Trizol and immediately homogenized using soft tissue homogenizing tubes (Precellys P000912-LYSK1-A.0, P002391-P24T0-A.0). RNA was purified from homogenized samples using the RNeasy Mini Kit (Qiagen 74106) according to manufacturer instructions.

### Knee hyperalgesia assessment

Knee hyperalgesia was assessed in a blinded fashion using an algesiometer (IITC Life Science, Woodland Hills, CA), as previously described^40^. Briefly, the tip of the applicator was positioned on the medial aspect of the knee joint, and force was gradually applied until a nocifensive response occurred (struggle or vocalization), which was recorded as the withdrawal threshold. An average of triplicate threshold measurements was calculated for each joint.

### Human adipose spheroids and glycerol assay

Subcutaneous human preadipocytes (ZenBio Cat#ZBM001.08 Lot#LM042624B) were seeded at a concentration of 12.5k cells per well in a Nunclon Sphera 96wp ULA U-bottom plate (#174925 Thermo Fisher) in maintenance media containing αMEM (Sigma Aldrich #M4526-500ML) supplemented with 1% GlutaMAX, 1% Penicillin/Streptomycin and 10% FBS (Merck #C8056-500ML) for 3 days. After 3 days media was carefully removed and the formed spheroids were individually added to 30 µL of ice cold Matrigel® Matrix for Organoid Culture (Corning #356255) and plated on 12 well non adherent well plates, one spheroid per well. The plates were flipped and incubated for 30 min at 37C until Matrigel solidified. When Matrigel was solid, initiation media was added containing αMEM supplemented with 20% FBS, 5 µg/mL Insulin (Merck #I2643-50MG), 450 µM 3-isobutyl-1-methylxanthine (IBMX) (Merck #I5879-100MG), 1 µM Dexamethasone (Merck #D0700000), 2 µM Rosiglitazone (Merck #PHR2932-300MG) and 60 µM Indomethacin (Merck #I7378-5G) with or without 10mM Yoda1 (Sigma Aldrich #SML1558-5MG, LOT0000395581) and 10 ng/mL TGFβ (PeproTech, Cat#100-21-10UG, LOT#0724209) for 3 days. After 3 days, initiation media was replaced with Subcutaneous Preadipocyte Differentiation Medium (ZenBio #DM-2) with or without 10mM Yoda1 (Sigma Aldrich #SML1558-5MG, LOT0000395581) and 10 ng/mL TGFβ (PeproTech, Cat#100-21-10UG, LOT#0724209) for 14 days. Every 3-4 days media was changed. On day 14 after switching to differentiation media, media was collected and analysed with the Glycerol Assay Kit (Sigma-Aldrich #MAK117) according to manufacturers’ instructions. The organoids were fixed with 4% PFA for 30 minutes, washed with PBS and stained with 1:1000 BODIPY and 1:1000 DAPI Solution (BD Pharmingen™ #564907) for 1 hour. Following a PBS wash the organoids were imaged with an Andor BC43 Benchtop Spinning Disk microscope and processed with Fiji.

### Bulk RNA-sequencing

#### Sample quality assessment and submission

Phenol chloroform extraction followed by RNA isolation using a Qiagen RNeasy Mini Kit with DNase digestion was performed to isolate RNA. RNA quality was evaluated at the University of Michigan Advanced Genomics Core and by Novogene Corporation Inc (Sacramento, CA, USA) **(Supplemental Tables 8 and 10)**. High quality samples (RIN > 5.0) were submitted for library preparation (poly-A enrichment) and paired-end sequencing using NovaSeq X Plus. HISAT2 (v2.0.5) was used to align raw FASTQ files to the GRCm38/mm10 genome. FeatureCounts (1.5.0-p3) was used to generate count matrices based on reads mapped to each gene.

#### Differential expression and pathway analysis

Stringent quality control was performed to preserve only high-quality genes (FPKM > 1 and total reads > 10). Sex-linked (*Xist*, *Uty*, *Eif2s3y*, *Kdm5d*) and non-protein-coding genes were also excluded. pcaExplorer (v.3.4.0) was used to perform principal component analysis (PCA); no samples were identified as outliers based on a 95% confidence interval. DESeq2 (v1.50.2) was used for differential expression analysis (Wald test, Benjamini-Hochberg correction). Gene lists were filtered (padj < 0.05, |log_2_FC| > 0.585) to produce significant differentially expressed genes (DEGs). To identify DEGs for the Yoda1 intra-articular injection experiment, paired statistics were used to compare ipsilateral and contralateral limbs from each mouse. For the mechanical loading experiment, unpaired statistics were used.

Pathway analysis was performed on PantherDB^91^ using the gene ontology complete biological processes database (GO:BP). All significant DEGs and their Log_2_FC values were submitted for statistical enrichment testing. Significant DEG lists were separated by positive or negative fold changes and submitted for statistical overrepresentation testing. Findings were corroborated by submitting all genes for gene set enrichment analysis (GSEA) using clusterProfiler^92^ (v.4.12.0). Positively enriched, significant DEGs were submitted to RcisTarget for transcription factor motif enrichment analysis using the default arguments for the built-in cisTarget() function. Significant, high-confidence TF motifs (NES > 3) with minimum average read counts of 0.1 were considered. TRUUST analysis was also performed on these DEG groups using Metascape^93^.

Publicly available bulk RNA-seq data of murine Sham, 7d and 28d ACLR whole synovium (n=12 mice per condition. 6 male, 6 female) from a prior publication by our group was downloaded from GEO (GSE271903)^40^. Differentially expressed genes and pathways were obtained as described in the original paper.

### Single cell RNA-sequencing

#### Human synovium

Synovial single cell RNA-seq data from healthy and OA human synovium were gathered from five publicly available datasets (GSE283080, GSE152805, GSE246416, GSE233502, GSE126651). Individual samples in each dataset were filtered based on robust quality control metrics: nCount > 1800, nFeature < 200, ratio of mitochondrial genes > 10% were filtered out. Samples were integrated using the Seurat pipeline, with SCTransform normalization and RPCA integration. Linear and non-linear dimensionality reduction was performed using PCA and UMAP, respectively. The resultant integrated object was clustered at a dimension of 35 and resolution of 0.25 to resolve broad cell type clusters. Fibroblasts (PDGFRa-hi) were identified and subset, then re-clustered using a dimension of 25 and resolution of 0.25 resulting in 12 distinct clusters. FindAllMarkers() was used to find marker genes for each cluster (padj < 0.05). Pathway analysis using statistical overrepresentation testing in PantherDB^91^ of marker genes was used to identify the SF subtypes (Supplemental Table 5). Four lining clusters (PRG4+ VCAM+) and eight synovial sublining clusters were identified. scProportionTest was used to calculate which clusters were significantly disease enriched (FDR < 0.05, |Log_2_FC| > 0.585). FindMarkers was used to calculate Log_2_FC of genes.

#### Murine synovium

Single cell RNA-seq of murine sham, 7d and 28d ACLR synovia was collected from a previously published dataset from our group (GSE211584)^8^. Integration and clustering were performed as described in the original paper to reproduce the identical UMAP and clusters. The FindMarkers() function with default arguments was used to perform differential expression analysis between conditions. Pseudobulk gene expression was calcuated using AggregateExpression() and z-scores were calculated. Differentiation trajectory analysis was performed using Monocle3 (v 3.0.0)^42^. *Dpp4*+ progenitors, *Acta2*+ myofibroblast-like SFs and *Prg4*+ lining SFs were subset and preprocessed with 22 dimensions. Pseudotime was calculated along this trajectory, with the root node set in the *Dpp4+* cluster. Genes with correlations > 0.15 along pseudotime and average normalized reads > 0.1 were grouped by covariance into modules. Genes from modules enriched in the trajectory towards myofibroblast-like SFs (modules 3 and 5) were submitted to transcription factor binding motif (i.e. regulon) analysis using RcisTarget^43^. Significant, high-confidence TF motifs (NES > 3) with average read counts > 0.1 were considered. NES and log-normalized expression of predicted TFs were plotted.

### Spatial Transcriptomics

Sham, ACLR 7d, and ACLR 28d hindlimbs (n=2 male and n=2 female per condition) were collected from C57BL/6J mice and processed for spatial transcriptomics according to our published protocol^94^. In brief, mice were placed under deep isoflurane-induced anesthesia and perfused with Z-fix Zinc Formalin Fixative (Cancer Diagnostics; SKU: 175). Excess muscle was removed and limbs were fixed for an additional 48 h in Z-fix at 4 °C, then decalcified in 80% (v/v in PBS) Immunocal decalcification solution (Fisher Scientific; STL141432) for 16h while nutating at 4 °C. Following paraffin processing at the Michigan Integrative Musculoskeletal Health Core Center’s Structure, Composition, and Histology Core, samples were individually embedded with the medial side facing down, and excess paraffin was trimmed to improve RNA extraction efficiency. RNA was isolated from five 20 µm sections using the Qiagen RNeasy FFPE kit with double the reagent volume to ensure thorough removal of residual paraffin that could compromise RNA quality assessment. Extracted RNA was submitted to the University of Michigan Advanced Genomics Core, where quality control was performed using an Agilent Bioanalyzer to assess RNA concentration, RNA integrity number (RIN), and DV200 values (**Supplemental Table 14**). After passing the RNA quality assessment, the proximal femur, distal tibia, and further excess muscle were removed from the block to isolate individual knee joints. Six joints were then co-embedded in a single paraffin block.

Sagittal sections of the co-embedded joints were collected on Xenium slides under sterile conditions inside a dead air box and submitted to the University of Michigan Advanced Genomics Core for processing using the 10x Genomics Xenium 5K Prime Mouse Pan Tissue & Pathways panel, including an additional custom panel of 100 genes previously reported^94^. The optional Multimodal Cell Segmentation was included, containing antibodies marking the cell membrane, cytoplasmic RNA, cytoplasmic protein, and nuclei to aid in segmentation of single cells. Slides were then imaged and transcripts were detected using the Xenium Analyzer (v3.3.0.1). Hematoxylin and Eosin (H&E) staining followed transcript detection and processing. H&E images for each sample were imported into Xenium Explorer for alignment with the Xenium morphology image^94^.

Anterior and posterior synovia were manually outlined in Xenium Explorer (v4.1.1), and coordinates were extracted. Synovial masks were subset from the Xenium data using the extracted coordinates. Only high-quality gene transcripts (Q-value ≥ 20) were considered for downstream analysis. All synovial masks were integrated using SCTransform normalization and RPCA integration in Seurat (v5.3.0)^95^. Dimensionality reduction was done using PCA and UMAP. Semi-supervised clustering of cells was done using a dimension of 30 and resolution of 0.35. FindAllMarkers() was used to calculate differentially expressed genes within each cluster, which were used to annotate major synovial cell types (**Fig S1A-E**). Fibroblasts were identified, subset, then re-clustered (dimensions = 15, resolution = 0.35) to identify SF subsets. SFs were annotated based on marker genes and concordance with markers from the murine single cell RNA-seq atlas. Cluster annotations were exported from R and imported into Xenium Explorer for visualization of synovial fibroblast subsets and gene transcripts on H&E images.

### Bioinformatics Analysis and Visualization

All bioinformatics analyses were performed in R (v4.5.2) and RStudio (v 2025.9.2.418). Bubble plots were created using ggplot2 (v4.0.3). Heatmaps of z-scores and fold changes of genes were made using pheatmap (v1.0.13). UMAPs, FeaturePlots, and ViolinPlots were visualized using Seurat (v5.5.0). Bar graphs were made using GraphPad Prism 11 (v11.0.0). Xenium Explorer (v4.1.1) was used for cell and transcript overlay, and image export.

## Author Contributions

Conceptualization: ECF, KDH, AJK, TM

Methodology: ECF, AM, SYK, HXT, LH, AJK, TM

Validation: ECF, SYK, TM

Formal Analysis: ECF, SYK, LL, AJK

Investigation: ECF, AM, AK, EK, HS, MW, SYK, LL, MDN, GE, RRS, SCH, SZL, HXT, DMN, KB, SGN, AJK

Resources: LH, RLM, CRS, KDH, TM

Data Curation: ECF, SYK, LL, TM

Writing – Original Draft: ECF, TM

Writing – Review & Editing: ECF, AM, LL, MDN, RLM, CRS, KDH, AJK, TM

Visualization: ECF, AJK

Supervision: ECF, LH, RLM, CRS, KDH, AJK, TM

Project Administration: ECF, KDH, AJK, TM

Funding Acquisition: ECF, LH, CRS, KDH, AJK, TM

## Acknowledgements

We would like to thank the staff at the UMICH Advanced Genomics Core, the UMICH Flow Cytometry Core, the UMICH Orthopaedic Research Laboratories Histology Core, and the Unit for Laboratory Animal Medicine. We would also like to thank the Michele Lab at U-M for their provision of the Flexcell® FX-4000T Tension Plus System.

## Data Availability

Mined sequencing datasets are publicly available from NCBI GEO (GSE211584, GSE271903, GSE283080, GSE152805, GSE246416, GSE233502, GSE126651). Newly generated sequencing bulk RNA-sequencing and spatial transcriptomics datasets have been deposited on Zenodo. These datasets will be made available following peer review.

