## Supplemental Figures for "PIEZO1 mediates mechanosensing and myofibroblastic activation of synovial fibroblasts in post-traumatic osteoarthritis"

### Supplementary Materials

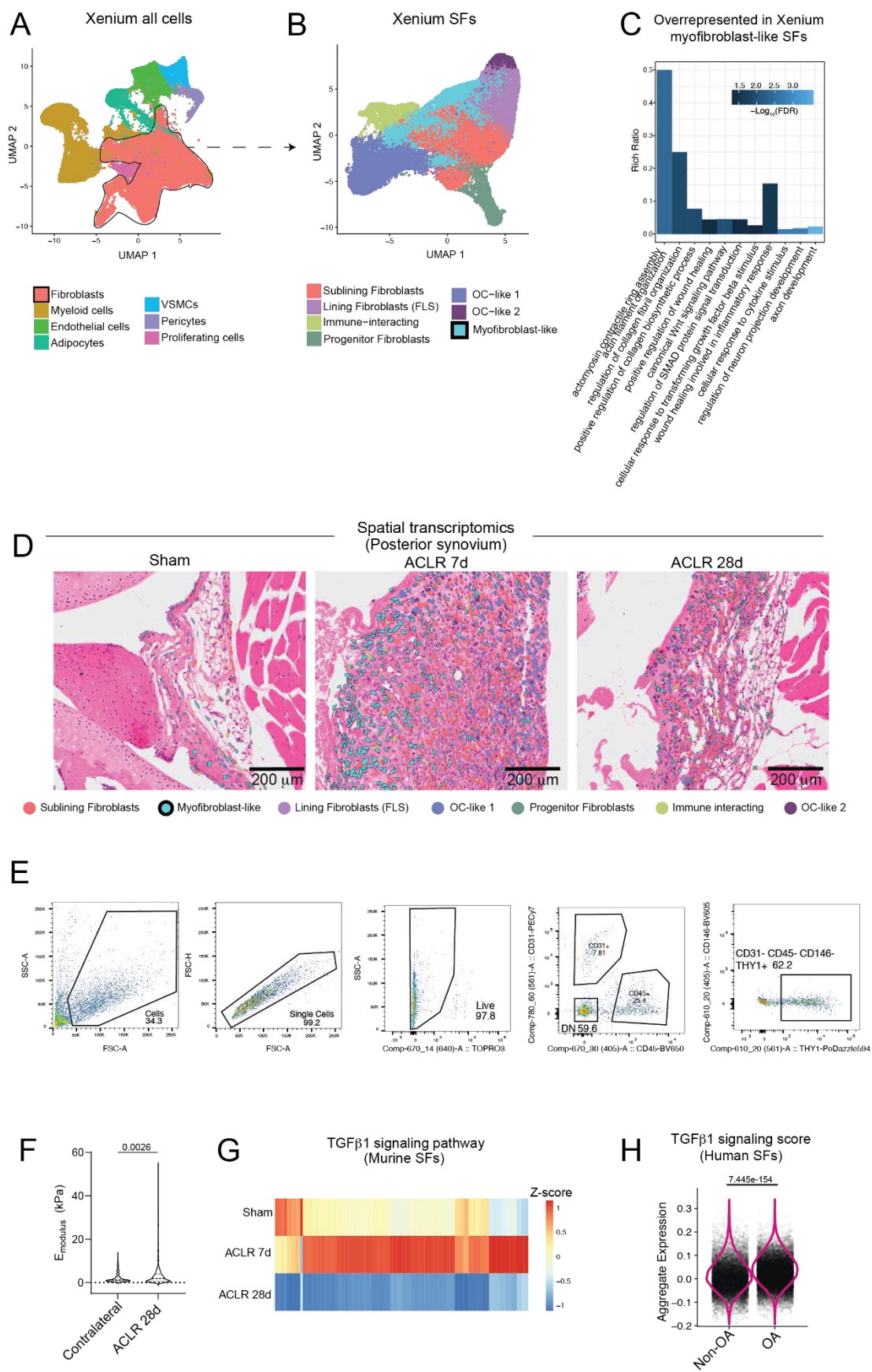

**Fig S1. (A)** UMAP plot of all cells identified by Xenium spatial transcriptomics. **(B)** UMAP plot of synovial fibroblast (SF) clusters identified by Xenium spatial transcriptomics. **(C)** Select myofibroblastic Gene Ontology – Biological Processes (GO:BP) terms significantly enriched in myofibroblast-like SFs identified by Xenium spatial transcriptomics. **(D)** Xenium

spatial transcriptomics showing spatial distribution of synovial fibroblast subsets in posterior synovium of Sham (right), ACLR 7d (middle), and ACLR 28d (right) mice. Representative H&E-stained sagittal sections with cluster overlays are shown. Myofibroblast-like SFs are outlined and highlighted in light blue. **(E)** Gating strategy for flow cytometry of *Acta2*-Cre<sup>ERT2</sup>;tdTom (*Acta2*;tdTom) synovium. Of live cells, CD31- CD45- CD146- THY1+ were identified as sublining fibroblasts, of which tdTom+ cells were quantified. **(F)** Violin plot of all indentation moduli (n=15-20 measurements/limb, 1 slide per mouse) measured in anterior synovium of contralateral and ACLR 28d mice. Unpaired two-tailed t-test with Welch's correction. **(G)** Relative expression by bulk RNA-seq of all genes comprising the GO:BP term "TGF-β1 signaling pathway" across Sham, ACLR 7d, and ACLR 28d murine synovium. Aggregate relative expression of n=12 per condition (6 male, 6 female) is plotted. Data sourced from, Lammlin et al<sup>40</sup>. **(H)** Violin plot (VlnPlot) of aggregate expression of all genes in the GO:BP term "TGF-β1 signaling pathway" in non-OA and OA human SFs, by single-cell RNAseq (Wilcoxin rank-sum test using FindAllMarkers).

A

| Accession | Author (publication year) | Number of samples | # SFs after QC | Represented sexes | Collection/surgery |
| --- | --- | --- | --- | --- | --- |
| GSE283080 | Miyahara (2025) | 4 OA samples | 8240 | Males | Total knee arthroplasty at end stage OA |
| GSE152805 | Chou (2020) | 3 OA samples | 7477 | 1 Male, 2 Females | Total knee replacement at end stage OA |
| GSE246416 | Smith (2024) | 5 Non-OA samples | 3597 | Males | Arthroscopy with needle or synovial biopsy for patients with no clinical sign of inflammation |
| GSE233502 | Faust (2024) | 3 Non-OA samples | 9413 | 8 Males, 8 Females (pooled) | Post-mortem from non-OA patient samples |
| GSE126651 | Tang (2019) | 3 OA, 3 Non-OA | 19491 | 4 Females, 2 Males | OA: total knee replacement at end stage OA<br>Non-OA: patients undergoing resection of bone tumors or from organ donors |

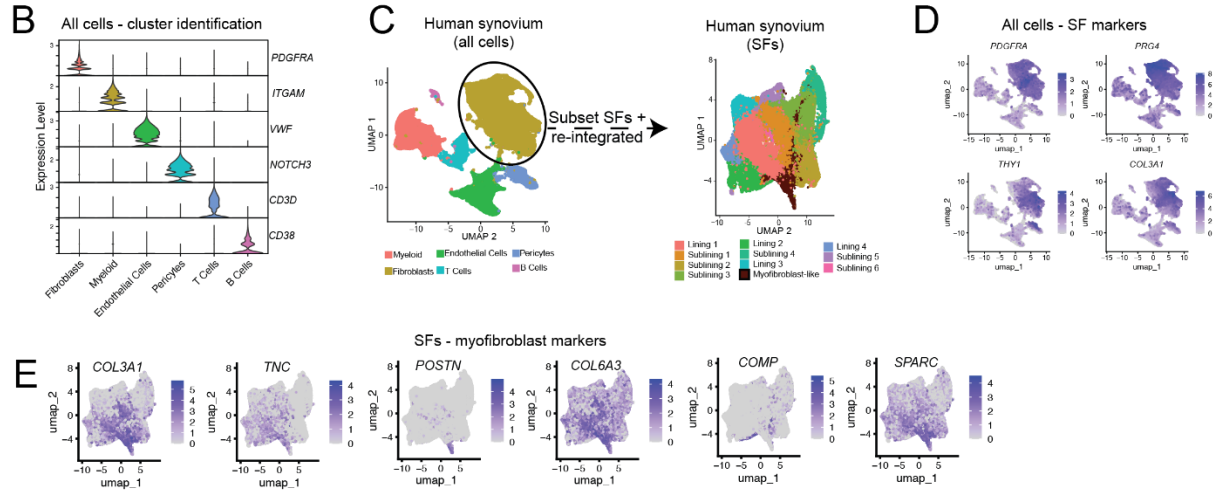

**Fig S2. A)** Demographic and quality control data of publicly available datasets used to generate the human SF atlas. **B)** Violin plot of marker expression of main synovial cell populations. **C)** Diagram of method used to generate SF clusters from human all cells data. **D)** Feature plots of markers used to identify SFs from all cells object. **E)** Feature plot of markers used to identify myofibroblasts from SF object.

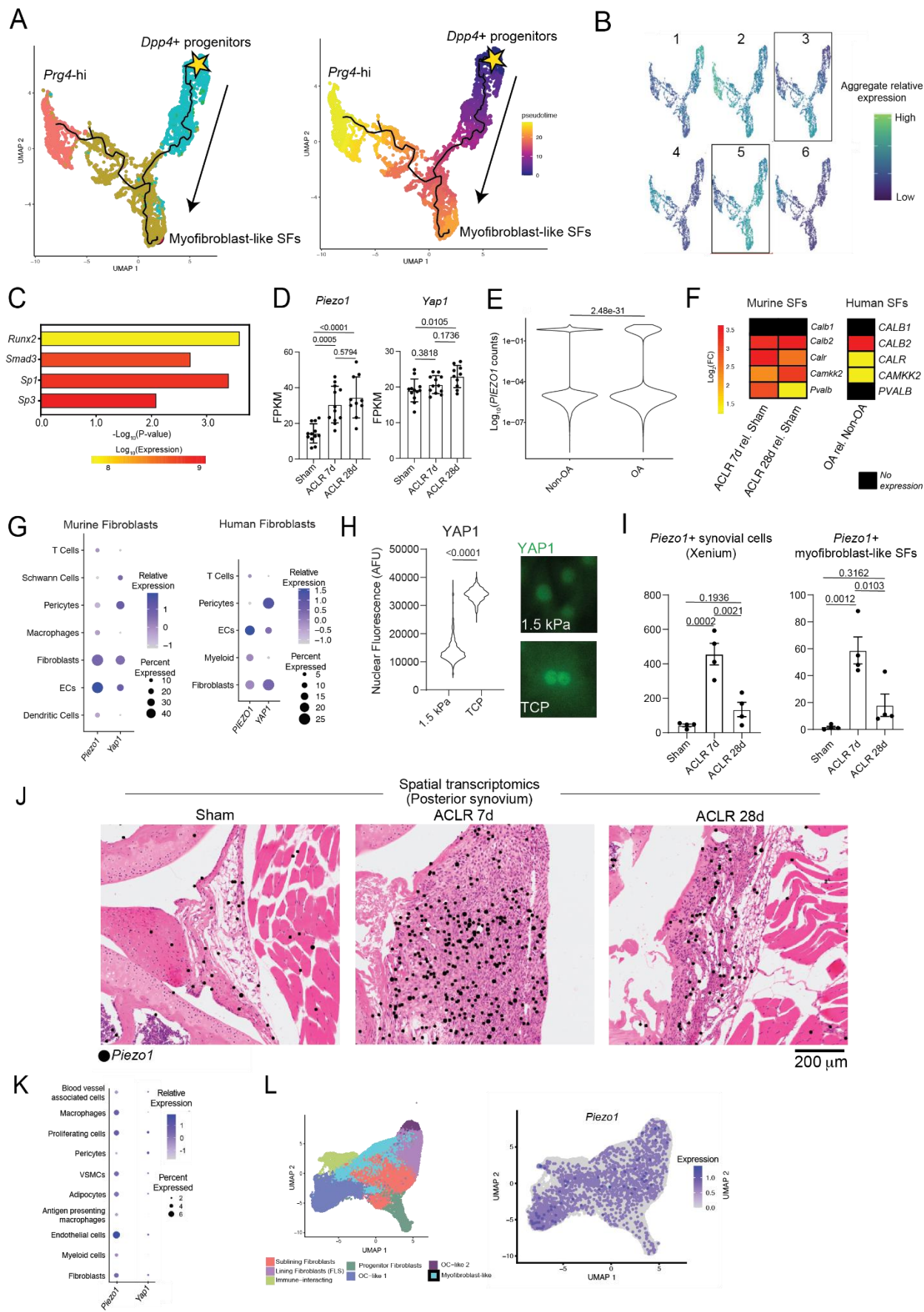

**Fig S3. (A)** UMAP plot of Monocle3-based cellular trajectory of murine *Dpp4*<sup>+</sup> progenitor-like SFs transitioning to either myfibroblast-like SFs or *Prg4*<sup>hi</sup> lining SFs, with cluster overlay (left) or pseudotime overlay (right). The bifurcation towards myfibroblast-like SFs was subset for Fig 2A. **(B)** UMAP plot of aggregate relative expression of pseudotime-dependent, co-variant genes modules identified by Monocle3. Modules 3 and 5 were enriched in the myfibroblast-like branch and used

for TF binding motif analysis. **(C)** Significantly enriched TFs associated with genes from Modules 3 and 5 identified by TRUUST analysis. **(D)** Gene expression, expressed in FPKM, of *Piezo1* and *Yap1* in Sham, ACLR 7d, and ACLR 28d murine synovium via bulk RNA-seq of whole synovium. Data sourced from Bergman+Lammlin et al<sup>40</sup>. n=12 per condition (6 male, 6 female). One-way ANOVA with Tukey post-hoc testing. **(E)** Log normalized *PIEZO1* SF counts in non-OA and OA synovium (Wilcoxin rank sum test). **(F)** Heatmap of Log2 fold-change of selected genes related to calcium ion shuttling in murine SFs in ACLR 7d and ACLR 28d relative to Sham (left) and in human OA SFs relative to non-OA SFs (right). **(G)** Dot plots of *Piezo1* and *Yap1* expression in major cell types in murine (left) and human (right) synovium. **(H)** Representative microscopy (right) and nuclear intensity quantification by arbitrary fluorescent units (AFU) (left) of YAP1 immunofluorescent staining of murine SFs cultured on soft substrate (1.5 kPa) or tissue culture plastic (TCP, ~1 GPa) substrate for 24h. Two-tailed independent t-test. **(I)** Number of all *Piezo1*+ synovial cells (left) and *Piezo1*+ myofibroblast-like SFs (right) detected by Xenium 5K spatial transcriptomics. N=4 mice (2 male, 2 female) per condition. One-way ANOVA with Tukey's post-hoc testing. **(J)** Xenium spatial transcriptomics showing spatial distribution of *Piezo1* transcripts (black dots) on representative H&E-stained sagittal sections of posterior synovium of Sham (left), ACLR 7d (middle), and ACLR 28d (right) mice. **(K)** Dot plot of *Piezo1* and *Yap1* expression across major synovial populations as detected by Xenium spatial transcriptomics. **(L)** Xenium spatial transcriptomics-based UMAP feature plot of *Piezo1* expression in SFs (right) next to UMAP plot of SF clusters (left, identical to Fig S1B).

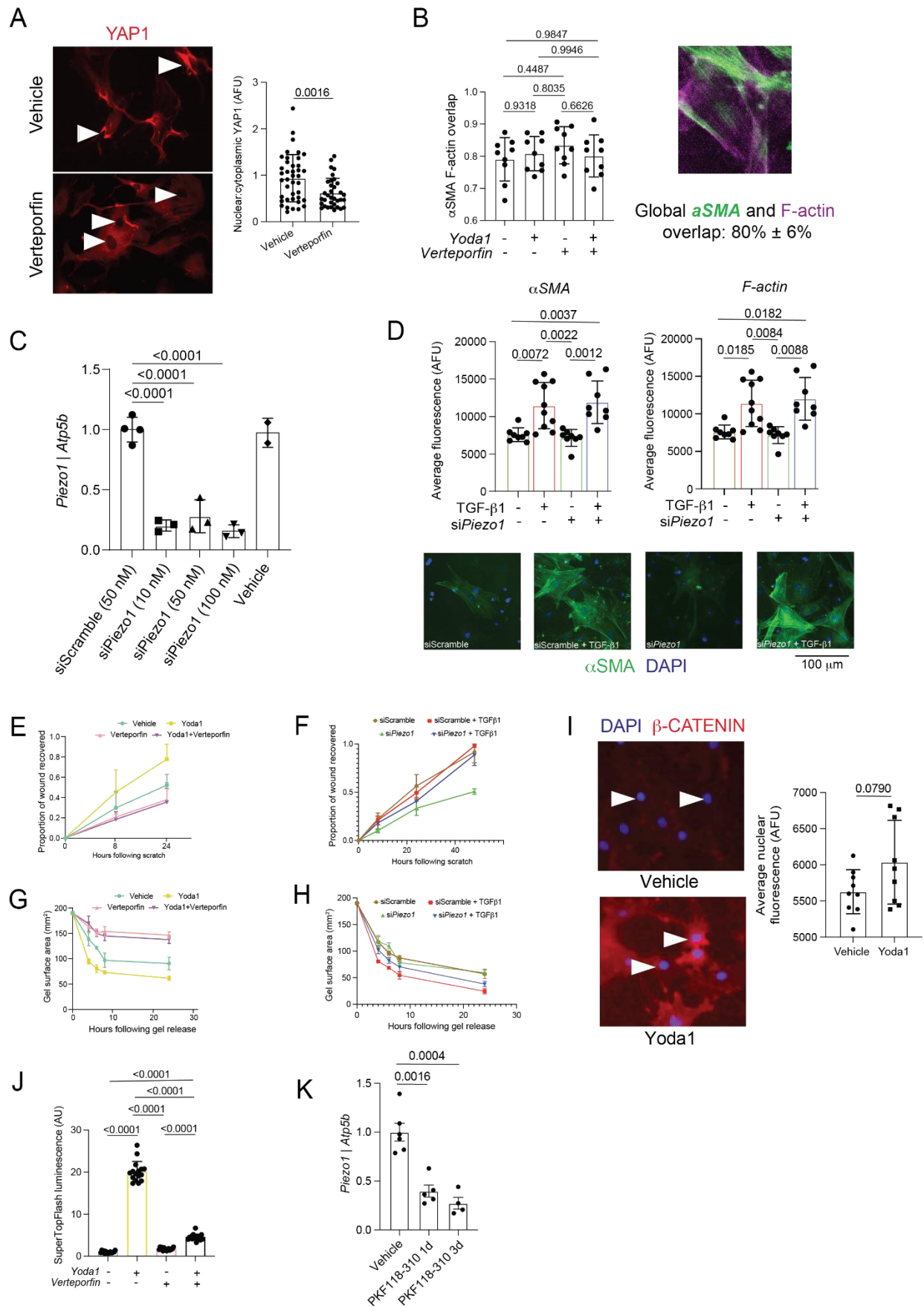

**Fig S4. (A)** Representative microscopy (left) and quantification of the ratio of nuclear fluorescence to cytosolic fluorescence (right) of YAP1 immunofluorescent staining of murine SFs treated with vehicle or Verteporfin for 24h. White arrows label nuclei. Replicates represent individual SFs. Two-tailed independent t-test. **(B)** Proportion of αSMA and F-actin overlap in SFs from experiment in Fig. 3A. n=3 technical replicate runs (each with n=6 biological replicates pooled), with three fields

of view quantified per replicate. One-way ANOVA with Tukey post-hoc testing. **C)** *Piezo1* gene expression by qPCR in murine SFs transfected with non-targeting siScramble, multiple si*Piezo1* concentrations, and vehicle. n=2-4 per condition. One-way ANOVA with Tukey post-hoc testing. **D)** Whole-cell fluorescent intensity quantification of αSMA (top left) and F-actin (top right) immunofluorescent staining of murine SFs pre-transfected with si*Piezo1* or siScramble followed by treatment with vehicle or TGF-β1 for 24h. n=8-10 technical replicate runs (n=6 biological replicates pooled) per condition. One-way ANOVA with Welch's correction and Tukey post-hoc testing. Representative fluorescent images of αSMA are shown on the bottom, with nuclei labeled by DAPI. **(E-F)** Time-dependent line plots of the proportion of scratch wound recovered by SFs in experiments from scratch wound assays in Fig. 3B and 3D, respectively. **(G-H)** Time-dependent line plots of collagen gel surface area from gel contraction experiments in Fig. 3C and E, respectively. **(I)** Representative microscopy (left) and nuclear intensity quantification (right) of active β-catenin immunofluorescent staining of murine SFs treated with vehicle or Yoda1 for 24h. n=3 technical replicates (n=6 biological replicates pooled), with three fields of view quantified per replicate. Two-tailed independent t-test. **J)** SuperTopFlash luminescence normalized to cell viability of HEK293-SuperTopFlash cells treated with vehicle, Yoda1, Verteporfin, or Yoda1 + Verteporfin for 24h. n=16 wells per condition. One-way ANOVA with Tukey post-hoc testing. **K)** *Piezo1* gene expression by qPCR in murine SFs treated with vehicle or the cWnt inhibitor PKF 118-310 for one or three days. n=4-6 technical replicates (n=6 biological replicates pooled). One-way ANOVA with Tukey post-hoc testing.

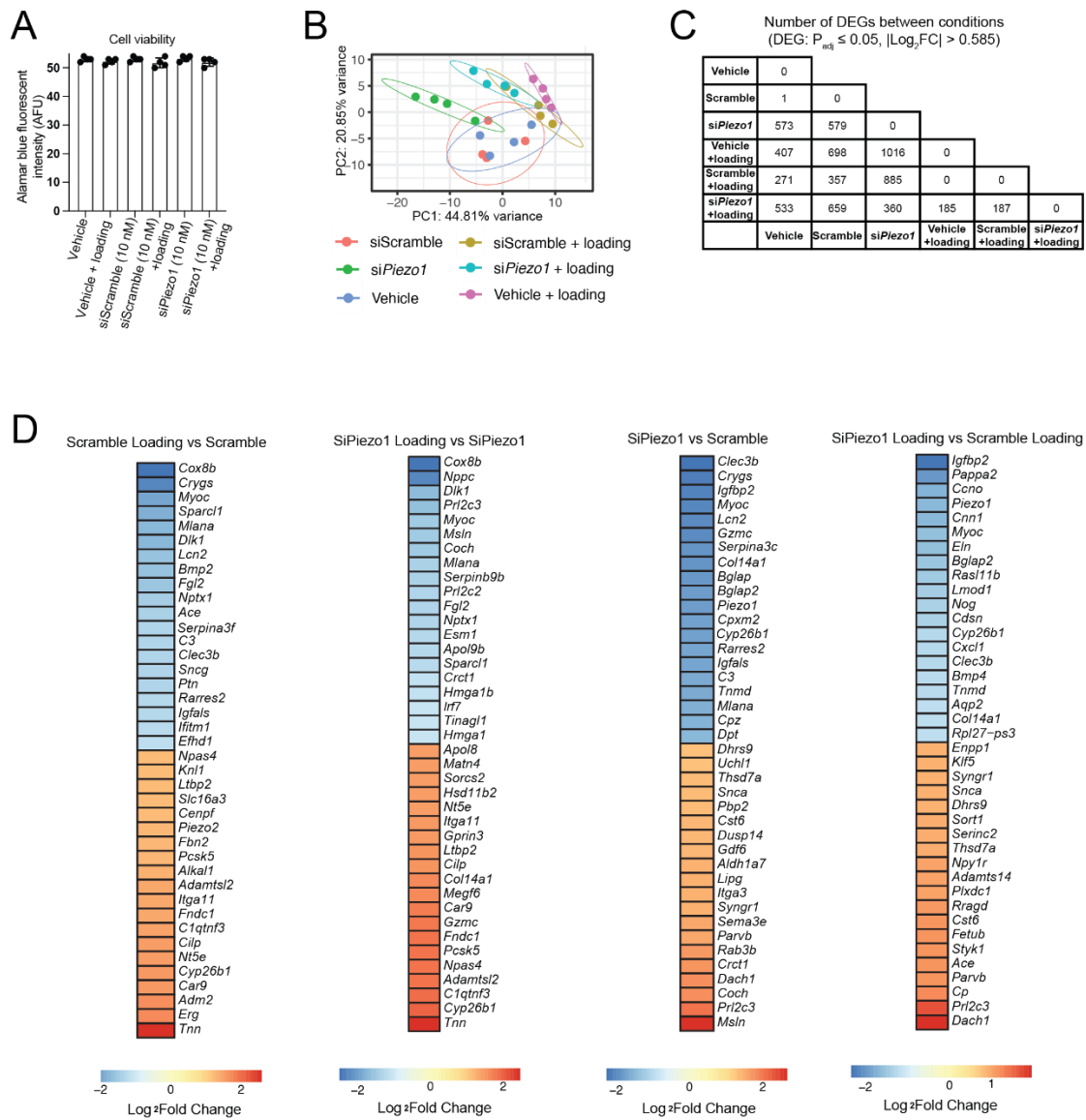

**Fig S5. A)** Alamar Blue fluorescence (AFU) to assess cell viability of SFs treated for 24h with siScramble and siPiezo1 under unloaded and loaded conditions. $n=6$ biological replicates (4 technical replicate runs). **(B)** Principal component analysis plot of bulk RNAseq of SFs from the mechanical loading experiment in Fig. 4. $n=4$ per condition. Ellipses represent the 95% confidence intervals. **(C)** Summary table of the number of DEGs ($p_{adj} < 0.05$, $|\text{Log}_2\text{FC}| > 0.585$) between all conditions in SF mechanical loading experiments. **(D)** Differential expression ranked by log<sub>2</sub> fold-change of the top 20 upregulated (red) and top 20 downregulated (blue) DEGs across relevant group comparisons.

A

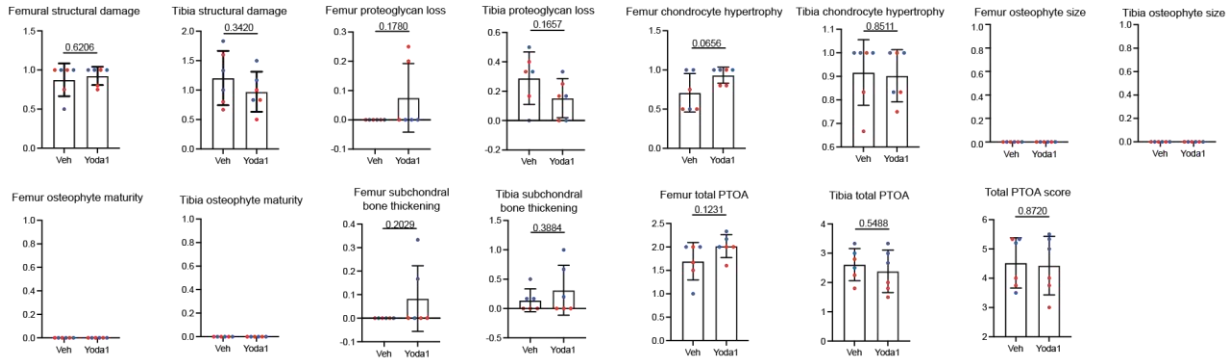

B

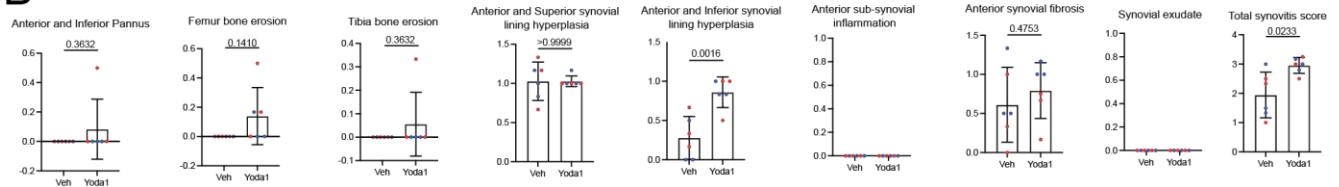

**Fig S6. (A-B)** Histological subscores and total scores for PTOA scoring (A) and synovitis scoring (B) of murine joints injected with vehicle or Yoda1. n=6 per condition (3 male, 3 female). Two-tailed paired t-tests, with Welch's correction applied to "total proteoglycan loss", "femur chondrocyte hypertrophy", "femur subchondral bone thickening", "Anterior and Inferior Pannus", "Femur bone erosion", "Tibia bone erosion", and "Total synovitis score" categories.

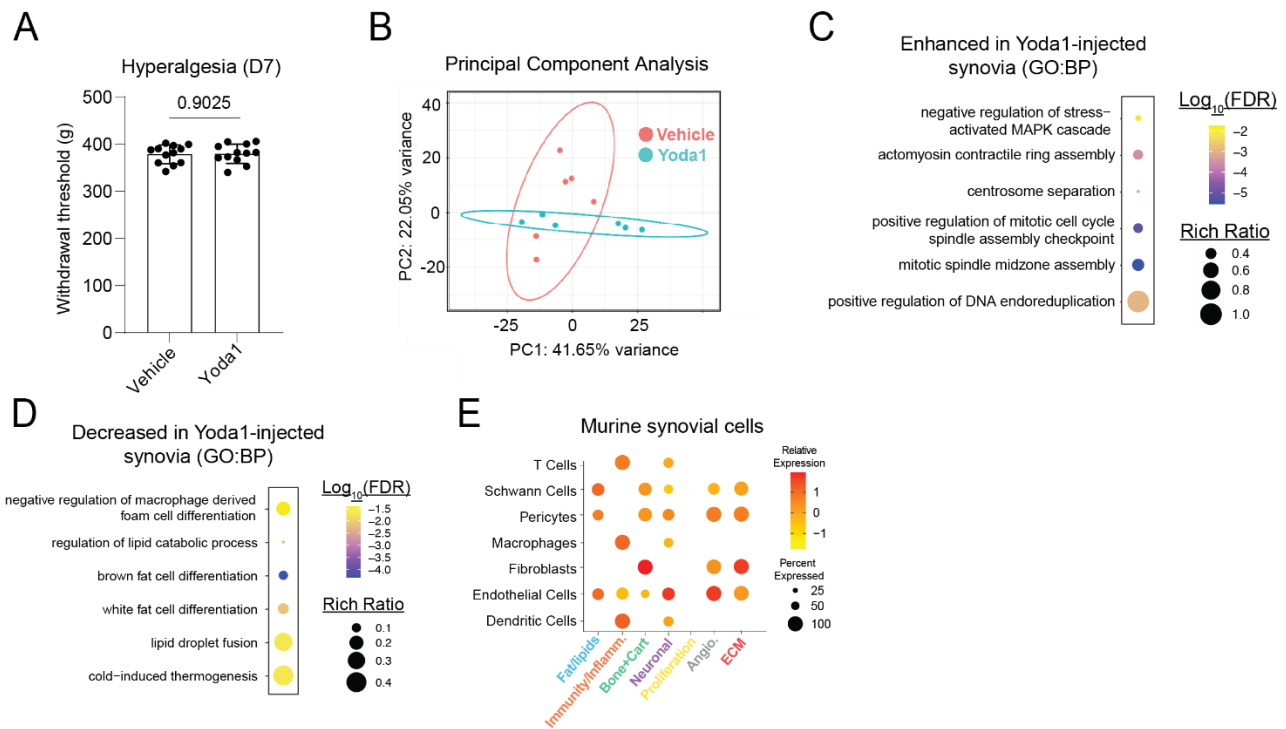

**Fig S7. (A)** Withdrawal threshold from blinded knee hyperalgesia assessments in joints administered Yoda1 or vehicle for five consecutive days. Measurements were taken two days after the last injection. $n=6$ mice per condition (3 male, 3 female). Two-tailed paired t-test). **(B)** Principal component analysis plot of bulk RNAseq of murine synovium from Yoda1-injected and vehicle-injected joints. $n=6$ mice per condition (3 male, 3 female). Ellipses represent 95% confidence intervals. **(C-D)** Selected significant ($\text{padj} < 0.05$) GO:BP pathway terms enhanced in (C) or downregulated in (D) synovium from Yoda1-injected vs vehicle-injected joints. Statistical overrepresentation of upregulated or downregulated DEGs, respectively. **(E)** Dot plot of aggregate expression of genes comprising select GSEA terms from Fig 5H across major murine synoviocyte subsets.

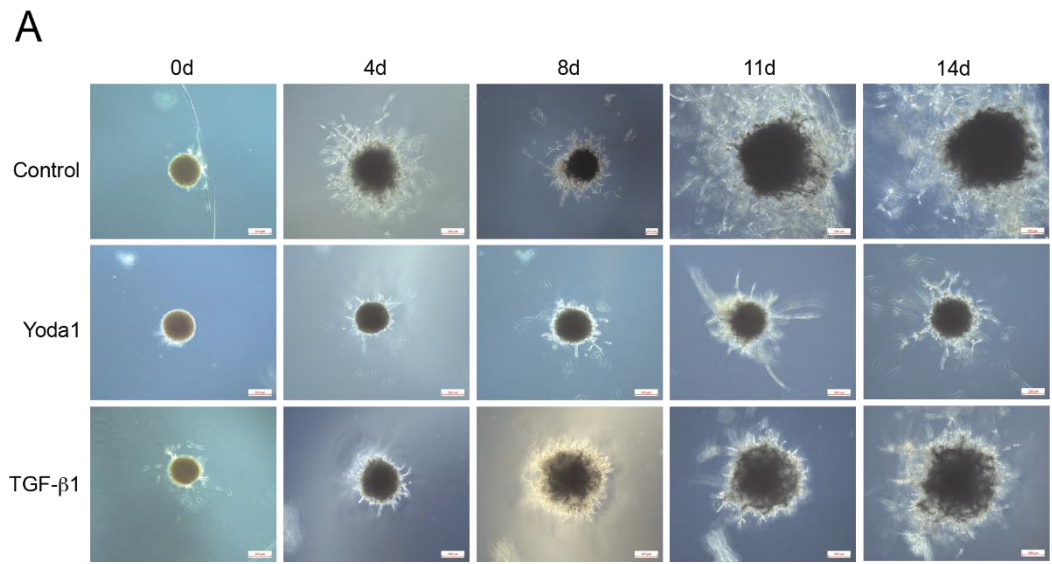

**Fig S8. (A)** Longitudinal brightfield confocal images of human preadipocyte organoids following induction of differentiation with pro-adipogenic media (ZenBio medium) with and without addition of Yoda1 (10 mM) or TGF-β1 (10 ng/mL).
